# Taxonomic Monograph of *Saxicolella* (Podostemaceae), African waterfall plants highly threatened by Hydro-Electric projects, with five new species

**DOI:** 10.1101/2021.06.19.449102

**Authors:** Martin Cheek, Denise Molmou, Sekou Magassouba, Jean-Paul Ghogue

## Abstract

The genus *Saxicolella* Engl. (Podostemaceae) are African rheophytes, restricted to rapids and waterfalls as are all members of the family. Previously, *Saxicolella sensu lato* was shown to be polyphyletic with two separate clades in the molecular phylogenetic study of Koi *et al*. (2012). The name *Pohliella* Engl. was recently resurrected for one clade that is sister to the American genera *Ceratolacis* (Tul.)Wedd., *Podostemum* Michx. and all Old World Podostemoideae (podostemoids) (Cheek 2020). *Pohliella* has distichous phyllotaxy, bilocular ovaries, filiform roots with paired holdfasts, and rootcaps. The second clade, *Saxicolella sensu stricto*, including the type of the generic name, has spiral phyllotaxy, unilocular ovaries, ribbon-like or crustose roots that lack both holdfasts and rootcaps. *Saxicolella sensu stricto*, sampled from the type species, *S. nana* Engl. of Cameroon, is embedded within and near the base of the major clade of African podostemoids and is sister to all other African genera apart from *Inversodicraea* R.E.Fr. and *Monandriella* Engl. Recently reduced to three species in Cameroon and S.E. Nigeria by the resurrection of *Pohliella* (3 – 4 species in Ghana and Nigeria-Cameroon), *Saxicolella* sensu stricto is expanded to eight species in this monograph by description of five new taxa. *Saxicolella futa* Cheek and *S. deniseae* Cheek are newly described from Guinea, *S. ijim* Cheek from Cameroon, the informally named *S*. sp. A from Gabon, and *S. angola* Cheek from Angola. The known geographic range of the genus is thus expanded c. 2,500 km westwards to Guinea from eastern Nigeria and c.1,500 km southeastwards from near Yaoundé to Cuanza do Sul, Angola. The greatest concentration of species occurs in the Cross-Sanaga interval of western Cameroon and eastern Nigeria, with three species. Cameroon (3 species) followed by Nigeria and Guinea (2 species each) are the countries with highest species diversity. The genus can be expected to be found in Sierra Leone, Liberia, Ivory Coast and Congo Republic. A classification is proposed grouping the species into three subgenera (*Saxicolella, Butumia* (G.Taylor) Cheek comb. et. stat. nov. and *Kinkonia* Cheek subgen. nov.) based on root morphology and shoot position and morphology.

The discovery, morphology, circumscription, distribution, and ecology of *Saxicolella* is reviewed, an identification key to the species is presented, together with descriptions, synonymy, links to illustrations, and extinction risk assessments for each of the eight species now recognised. All of the species are provisionally assessed as either Endangered or Critically Endangered using the IUCN 2012 standard, making this genus among the most threatened of its size globally. The major threats, above all, are hydro-electric projects. *Saxicolella deniseae* may already be globally extinct, and two of the four known locations of *S. angola* appear lost, *S*. sp. A of Gabon is threatened at at least one of its three locations, while *Saxicolella futa* is threatened at all three locations, all due to incipient or active hydro-electric projects. Contamination of watercourses by increased turbidity from silt-load due anthropic changes and by eutrophication from pollution are also threats for the majority of the species.

## Introduction

Podostemaceae are a pantropical family of annual or perennial herbs placed in Malpighiales in a sister relationship with Hypericaceae (Ruhfel *et al*. 2011). There are about 300 species globally, in c. 54 genera (Koi *et al*. 2012). Species numbers are highest in tropical America, followed by Asia, with Africa having c. 106 species (Cheek & Lebbie 2018). All species of the family are restricted to rocks in rapids and waterfalls of clear-water rivers (rheophytes) or occur in the spray zones of waterfalls (this paper). However, waterfalls are being increasingly exploited for hydropower at risk to the survival of the Podostemaceae they contain (Schenk *et al*. 2015; Cheek *et al*. 2015; Cheek & Ameka 2016; Cheek *et al*. 2017a; Cheek *et al*. 2017b). Most of the African species of Podostemaceae are narrow endemics, many being known from only a single waterfall. New discoveries of species are still made frequently, in addition to those studies above (Rial 2002; Cheek 2003; Schenk & Thomas 2004; Beentje, 2005; Cheek & Ameka 2008; Kita *et al*. 2008; Cheek & Haba 2016; Cheek *et al*. 2017b; Cheek *et al*. 2019a; Cheek *et al*. 2020a), including a new genus to science (Cheek & Lebbie 2018).

Three subfamilies are recognised. Tristichoideae, sister to the rest of the family, have three foliose tepals that protect the developing flower, and are tricarpellate. Weddellinioideae, with a single genus are Neotropical and have two foliose tepals and bilocular ovaries. Podostemoideae, pantropical, is the most genus- and species-rich subfamily. It has flowers protected in a spathellum, a balloon-like sac in which the flower develops while the plant is underwater, and tepals reduced to vestigial, filiform structures. African Podostemoideae, or podostemoids, are the main focus of this paper.

Important characters in defining genera in African podostemoids are the position of the flower in the unruptured spathellum, and the number of locules, shape, and sculpture of the ovary. At species level, important characters are the shape and relative proportions of spathellae, stigmas, anthers, filaments, gynophores, pedicels, and leaves.

The current taxonomic framework for African Podostemaceae was set in place by the revisions and Flora accounts of Cusset (Cusset 1973; Cusset 1974; Cusset 1978; Cusset 1983; Cusset 1984; Cusset 1987; Cusset 1997). Only recently has accumulating molecular phylogenetic data begun to influence the classification (Moline *et al*. 2007; Thiv *et al*. 2009; Schenk *et al*. 2015). Cusset’s work has been compiled and updated by Rutishauser *et al*. (2004) who recognise c. 85 species in 16 genera.

However, *Saxicolella* Engl. was one of the few African genera that Cusset did not revise. Yet, in her Flore du Cameroun account (Cusset, 1987), in addition to the type species *Saxicolella nana* Engl., she included in *Saxicolella* the genus *Pohliella* Engl. with two species *P. laciniata* Engl. and *P. flabellata* G. Tayl. Taylor (1953) had already expressed his doubts about *Pohliella* “Apart from differences in habit and shape of the stigmas, I am not satisfied that the key characters used by Engler to distinguish *Pohliella* from *Saxicolella* are sufficiently diagnostic.” The two genera were subsequently treated as synonymous under the name *Saxicolella* (e.g., Cook & Rutishauser 2001; Cook & Rutishauser 2007). It is not difficult to see why this was the case. Both genera are unusual among African Podostemoideae in that the ovary is not inverted in the spathellum but erect, and also in that at anthesis, the ovary is not held on a long pedicel that exceeds the length of the spathellum by 3 – 4 times but is either held inside the ruptured spathellum with only the styles and stamen emerging, or the pedicel is only as long or at most 2 times as long as the spathellum. Further, both genera have unistaminate flowers, which are not common in African Podostemoideae, where two stamens per flower are usual.

However, the molecular phylogeny of Koi *et al*. (2012) showed that *Saxicolella* in this broad sense was polyphyletic, with two clades arising at different points from the family tree. It was shown (Cheek 2020) that these two clades differ from each other in several important characters, sufficient to merit generic separation (Table 1 below, reproduced from Cheek 2020).

**Table 1.** Characters separating the two polyphyletic clades of *Saxicolella sensu lato* of Cook & Rutishauser (2001; 2007). Characters taken from Koi *et al*. (2012: 473, table 3), Ameka *et al*. (2002), Hall (1971), Engler (1926) and Cheek (pers. obs. in Ghana, 1995 and Cameroon, 2008). Reproduced from Cheek (2020). The Brazilian genus *Cipoia* with two species (Bove *et al*. 2006) is morphologically identical with *Pohliella* but molecular support is advised before synonymising it (Cheek 2020).

|  | *Pohliella* | *Saxicolella* |
| --- | --- | --- |
| Root caps | Present* | Absent |
| Phyllotaxy of main stems | Distichous** | Spiral |
| Root morphology | Subfiliform (slightly dorsiventrally flattened), <1 mm width | Broadly ribbon-like (width >10 mm except *S. futa*) or crustose |
| Paired root holdfasts exogenous | Present | Absent |
| Ovary | Bilocular | Unilocular |

\* recorded in *Pohliella submersa* Cheek (Ameka *et al.* 2002).
\*\* in *Pohliella laciniata* while the main stem has distichous phyllotaxy, the leaves subtending the terminal inflorescence are spirally arranged.

Therefore, *Pohliella* was proposed for resurrection (Cheek 2020), leaving *Saxicolella* in the narrow sense, with two species in Cameroon, one of which extended to Nigeria and one endemic to Nigeria. In this paper we provide a taxonomic revision of the genus *Saxicolella* including both newly collected and previously overlooked material that conforms to the delimitation of the genus as represented in Table 1. We also review what information is available about the genus.

## Material and methods

Four of the eight species accepted in this paper have been studied in the wild by the authors. Fieldwork to collect data on *Saxicolella* for this paper began as part of general botanical surveys in Cameroon for conservation management. The methodology used was as reported in Cheek & Cable (1997), and specimen data storage by Gosline, in Cheek *et al*. (2004). Fieldwork on the genus recontinued in Guinea nearly 20 years later as part of targeted surveys partly focussed on waterfalls, where Google Earth was used to target and navigate to waterfalls. A blade was used to remove plants from their rocks when exposed in the dry season, rehydrating them first if already desiccated. Conventional herbarium and silical gel specimens were made and photos were also taken where possible. The most complete set of material was deposited in the National Herbarium of the country concerned.

Herbarium material was examined with a Leica Wild M8 dissecting binocular microscope. This was fitted with an eyepiece graticule measuring in units of 0.025 mm at maximum magnification. Botanical line-drawings were made using the same equipment, fitted with a camera lucida. The morphological species concept was followed in defining species (each species being separated from its congeners by several, usually qualitative, morphological disjunctions), and the overall morphology of species was described and illustrated based on herbarium specimens following standard botanical procedures as documented in Davis & Heywood (1963). All specimens cited have been seen by the first author unless indicated ‘n.v.’ Herbarium citations follow Index Herbariorum (Thiers *et al*., continuously updated) and authors of plant names IPNI (continuously updated). Material or images were studied from, and checks made for specimens at B, BM, COI, EA, GC, HNG, K, L, LISC, MO, P, SCA and YA. Key online specimen databases searched included GBIF.org and Tropicos.org (continuously updated), MNHN collections website (continuously updated) and those of COI: https://coicatalogue.uc.pt/?Collector=gossweiler&t=results&orderby=relevance&orderdirection=D_ESC&size=10 and LISC: https://actd.iict.pt/list/?cat=quick_filter&search_keys

We were not able to inspect the DNA voucher specimen listed by Koi *et al*. (2012) as at TNS, but this is thought to be from the type location where standard herbarium specimens have already been made, and have been viewed. In total 16 unique herbarium records of the genus were studied not counting their duplicates. Nomenclatural changes were made according to the Code (Turland *et al*., 2018).

Conservation assessments were either taken from the recent literature (see citations) or made using the categories and criteria of IUCN (2012). The cell-size used for calculating area of occupancy was 4 km^2^, as advocated by IUCN. Geocat (Bachman *et al*. 2011) was used to calculate extent of occurrence in the two species with more than two locations.

## Results

### Discovery

The first published and type species of *Saxicolella, S. nana*, was collected in Kamerun, then a German colony, now Cameroon, in January 1914 by the renowned botanist Mildbraed (Engler 1926). In 1922 Gossweiler in Angola first collected material of the species published in this paper as *Saxicolella angola* Cheek. Keay, collecting in eastern Nigeria, in 1948 and 1950 respectively made the specimens that became the basis of *Saxicolella flabellata* (G. Taylor) Cheek (originally published as *Pohliella)*, and *Saxicolella marginalis* (G. Taylor) Cheek (originally published as the monotypic *Butumia*). In 1998 the first author collected in Cameroon and misidentified as *Ledermaniella* cf. *musciformis* the species published in this paper as *Saxicolella ijim* Cheek. Then, in Guinea-Conakry in Jan. 2018 he collected the material of the species named here as *Saxicolella futa* Cheek sp. nov., together with the second author. The second and third authors then went on in Feb. 2018 to collect the species described as *Saxicolella deniseae* Cheek *sp*.*nov. Saxicolella* sp. A only came to our attention as this paper was being concluded in mid-2021, thanks to photos on Tropicos via GBIF.org of recent collections by the LBV-MO botanical team.

The new species published in this paper are unlikely to be the last added to the genus. It is expected that botanical survey of the many rapids and waterfalls of Africa that have never been inspected for Podostemaceae will produce additional species new to science if this can be done before they are modified by the hydro-electric projects which are likely to result in their extinction.

### Morphology

While species of several other genera of African Podostemaceae have been investigated in detail for their morphology and anatomy in such studies as Moline *et al*. (2007) and Thiv *et al*. (2009), this has not been the case for any of the species of the genus *Saxicolella* Engl. as delimited here (the Ghanaian species previously referred to as *Saxicolella* have been transferred to *Pohliella*, see Cheek 2020). None of the species appear to have been investigated anatomically, nor has their micromorphology been investigated under the electron microscope. The overview presented here is partly based on the protologues of the species already published by Engler (1926) and Taylor (1953), but mainly from the observations of the authors of the four new species described below.

#### Root

The root (thallus) is either crustose-discoid and/or with several, ± broadly ribbon-like arms radiating from a central crustose area (the central crustose area is rarely absent/not detected e.g., *Saxicolella futa*). It is usually several times wider than thick and is closely appressed to the substrate of smooth rock to which is firmly fastened by numerous short root hairs. A faint raised ridge running along the midline of the root-ribbon of *S. futa* suggests that as in *Inversodicraea* (Cheek *et al*. 2017b), a single, central vascular bundle is present. Photosynthesis seems to be mainly performed by the ribbon-like roots since these make up most of the surface area of the plants, in fact >90% of the surface area in almost all species. Root-caps have not been reported nor observed but are in any case not usual in those Podostemaceae genera with crustose and broad ribbon-like roots. Nor are haptera, also known as hold-fasts, present. Roots are neither described nor preserved in the available material of *Saxicolella angola* and are incompletely known in *S. nana* and *S. flabellata*. The ribbon-like roots of individuals appear to radiate out from the central point of establishment, presumably where a seed has germinated and established. In contrast, in *Saxicolella nana* the radial growth appears to be “crustose”, that is, not to in the form of distinct separate ribbon-like structures, but a solid mass which extends outwards more or less evenly along the circumference, with only slight lobing at the margins.

In most species, e.g., *Saxicolella deniseae*, and *S. marginalis* the root is intermediate between the two extremes since it has both a central crustose part several centimetres diameter, but also the margins are well-developed into radiating ribbon-roots. In *Saxicolella futa* the central crustose part if developed at all, must be small and only a short-lived stage which is lost by fruiting time (if it is developed in the first place), leaving the radiating roots disconnected from each other at the centre.

In most species the ribbon-like, radiating roots rarely (*S. deniseae, S. marginalis* and *S. ijim*) branch, when they bifurcate into two equal branches. However, in *Saxicolella futa* the branching is frequent and regular, and the roots form a distinctive pattern. In fact, each species of *Saxicolella* can be identified by the architecture and gross morphology of its root alone (where known), although this can be difficult to convey in words.

#### Shoots

The origins of the shoots from the roots and their development, follows one of three patterns:

1. the shoots arise only from the central, more or less disc-like, crustose part of the root, and not from the radiating ribbon-like roots. The shoots form visible stems with measurable internodes. *Saxicolella nana, S. flabellata, S. ijim* and, possibly, (root unknown but visible stems present) *S. angola*.
2. The shoots arise only from the margins of the radiating, ribbon-like roots. The shoots are sessile, not forming visible stems but supporting an inconspicuous rosette of reduced leaves and a terminal spathellum. *Saxicolella marginalis, S. deniseae, S*. sp. A
3. The shoots arise only from the synusiae of the bifurcations of the radiating, ribbon-like roots. As in 2, the shoots are sessile, not forming visible stems. *Saxicolella futa*.

These three shoot position patterns appear to have value in supporting generic delimitation in Asian podostemoids, (Koi *et al*. 2012:475), pattern 1=“D (dorsal surface of root)”; pattern 2=“P (*Paracladopus*-type)”; pattern 3=“C (*Cladopus*-type)”; (Koi *et al*. 2012).

The shoot patterns appear to correlate with the three root patterns (see Roots, above). The taxonomic significance is discussed below.

In those species where visible stems are developed, they are erect, terete, and in those species where they exceed more than 5 mm long, sparingly branched. In *Saxicolella ijim*, the stems are robust and free-standing at anthesis. This species was found in the spray zone of a waterfall (Cheek pers. obs.) and is not supported by water as appears to be the case of the more laxly stemmed *S. flabellata* which has the longest (21cm) stems in the genus, described as flowing in the protologue (Taylor 1953).

#### Leaves

The phyllotaxy is consistently spiral. The leaves are best developed in the species with pattern 1 shoot position, where visible stems are developed. The largest leaves are those of *S. flabellata* which are flabellate (dorsiventrally flattened with radiating lobes) and up to 3 cm long, 2 cm wide. Each leaf bifurcates or trifurcates up to four times, the ultimate segments are capillary. The base is sheathing. Stipules are inconspicuous.

Leaves in *S. angola* are poorly preserved, smaller, but otherwise similar, with fewer bifurcations and with stipules conspicuous. In *S. nana* the leaves are filiform-capillary and trifurcate, while those of *S. ijim* are unbranched and laterally compressed.

In pattern 2 species, *S. marginalis* and *S*.*deniseae*, whilst the shoots are reduced and visible stems are not formed, the leaves appear reduced to the sheathing, stipulate base with only a rudimentary blade, while in *S*. sp. A the linear blade is as long as the flower

In pattern 3 *Saxicolella futa* the leaves are reduced further than in pattern 2, to inconspicuous, minute 0.3 mm long concave sheaths with stipules and blade not developed.

Leaves of the type usual in African podostemoids are absent, or at least have not been observed– that is, those which are filiform, terete and bifurcate repeatedly in the distal half, and which are shed before anthesis.

#### Inflorescences

In all species flowers occur singly at the apex of shoots except in *Saxicolella flabellata* and *S. angola* where they are in terminal clusters. The developing spathellae are protected by the subtending leaves in the earliest stages. In fact, the leaves appear to function primarily as protective bracts in most of the remaining species of the genus. The spathellum varies from globose (*S. ijim*) to narrowly ellipsoid, sometimes with a small apiculus. It lacks a distinct stipe.

**The flower** is erect and held within the opened spathellum at anthesis. Generally, only the styles and anthers are exserted from the ruptured spathellum but sometimes all or part of the ovary is projected from the spathellum. However, in *Saxicolella nana* and *S*. sp. A. the ovary can be projected on a naked pedicel as long as itself. A short pedicel, two filiform (rarely spatulate) tepals that flank the single stamen, and a short gynophore are present (absent in *Saxicolella* sp. A), all concealed within the ruptured spathellum at anthesis. The anther-thecae often face away from each other (latrorse). Pollen is dyad (where available for study).

The ovary is either ellipsoid, e.g. *S. ijim, S. futa, S. marginalis*, or narrowly ellipsoid (*S. flabellata, S. deniseae, S. nana, S. angola, S*. sp.A). In the fruit there are eight longitudinal ribs extending from base to apex (*S. flabellata, S. marginalis, S. deniseae*, sp. A) or the commissural ribs are not developed, when only 6 ribs are developed (*S. angola, S. nana, S. futa, S*.*ijim*).

The two stigmas are filiform or narrowly botuliform (*S. nana, S*. sp. A, *S. angola, S. flabellata, S. deniseae*) or they are complanate (flat) and about as broad as long (*S. futa, S. marginalis, S. ijim*). The free-central axile placenta in the unilocular ovary is either narrowly spindle-shaped e.g. *S. angola, S. nana, S. futa* or broadly so, occupying about half the radius of the locular cavity in *S. ijim*. The seeds are all ellipsoid, completely covering the placenta, where known.

The fruit, as the ovary, is fully erect, and isolobous (the two valves are equal). The fruit is carried further out of the spathellum remains by the extension of the pedicel post-anthesis. The two valves dehisce but usually persist in the fruit. The seeds are mucilaginous where known as usual in the family.

## TAXONOMY

### Saxicolella

Engl. (Engler 1926:456), non J.B. Hall (1971: 133); non Ameka *et al*. (2002). Type species: *Saxicolella nana* Engl.

*Pohliella sensu* Taylor *quoad P. flabellata* (Taylor 1953:53) Heterotypic synonym

#### Rheophytic herbs

*Roots* ribbon-like and/or disc-like, crustose, highly dorsiventrally flattened, to at least five times as wide as thick, adhering to the substrate by root hairs on the ventral surface, rootcaps and haptera absent. *Shoots* erect, minute and supporting sessile leaf rosettes, the stem then not visible, then arising from the margins of the radiating ribbon-like part of the root or, (*S. futa*) the sinuses of the bifurcations of the ribbon-like root OR up to several cm long, unbranched or branched, arising from either the centre of the disc-like crustose part of the root. *Leaves* with spiral phyllotaxy, bases sheathing where known, with one or a pair of acute basal stipules in leaves subtending spathellae (stipules absent in *S. futa* and often in proximal leaves on a stem of other species), blades filiform or flattened, usually entire, rarely bifurcate (or trifurcate) in *S. flabellata* and *S. nana* respectively, blades reduced or rudimentary in *S. deniseae* and *S. marginalis* or absent in *S. futa. Flowers* single, terminal on shoots, rarely in clusters on main stem (*S. angola, S. flabellata*). Spathellae ellipsoid, sessile, rarely globose (*S. ijim*), apex often with mucro. Flowers erect in intact spathellum. Held completely partly within the opened spathellum at anthesis except in *S. nana* and *S*. sp. A. Pedicel accrescent in fruit. Tepals 2, filiform rarely spatulate (sometimes spatulate in *S. ijim*), flanking the stamen. Stamen single, exceeding ovary, thecae often divergent. Pollen in dyads. Gynophore present (except *S*. sp. A and *S. ijim*). Ovary unilocular, ellipsoid, not laterally compressed, isolobous, erect, 6 or 8-ribbed in fruit, ovules numerous around a columnar axil placenta, septum not detected. Stigmas 2, botuliform to filiform or complanate. *Fruit* dehiscing into two equal, persistent valves. *Seeds* ellipsoid, mucilaginous.

**DISTRIBUTION**. Tropical West Africa: Guinea, Nigeria, Cameroon, Gabon and Angola. Eight species. *Saxicolella* species are restricted to Africa and extend from the Guinea Highlands of Guinea-Conakry in west Africa (newly recorded here) to Angola (newly reported here) in western south-central Africa. They are not recorded from the Congo basin, nor eastern Africa. The geographic range of the genera *Talbotiella* Baker f., (Leguminosae, nine species of evergreen tree) recently also extended to Guinea (van der Burgt *et al*. 2018), is similar to that of *Saxicolella* although that genus does not extend to Angola (Mackinder *et al*. 2010). *Mischogyne* Exell (Annonaceae trees, five species, Gosline *et al*. 2019) also has a similar distribution but has an outlying species in Tanzania and one in DRC. The highest species diversity of *Saxicolella* is the Cross-Sanaga River interval of eastern Nigeria-western Cameroon which has three of the eight species: *S. marginalis, S. flabellata*, and *S. ijim*. The Cross-Sanaga River interval area contains the highest species and generic diversity of flowering plants per degree square in Tropical Africa according to several studies (Barthlott *et al*. 1996; Dagallier *et al*. 2020) possibly in part because it corresponds with the Cameroon Highlands (Cheek *et al*. 2001). Many of the species and some genera (e.g. *Medusandra* Brenan (Peridiscaceae, Breteler *et al*. 2015; Soltis *et al*. 2007) are both endemic and threatened. *Saxicolella* species are known only from the five countries mentioned but are likely to be found in intervening areas such as Sierra Leone, Liberia, Ivory Coast and Congo-Brazzaville. Of the eight known species, four are point endemics,

**HABITAT**. *Saxicolella* only grows, as with most Podostemaceae, in sites with seasonally or permanently, fast-flowing, well aerated, silt-free fresh water. They are always associated with waterfalls and rapids. Unusually among African Podostemaceae genera, about half the known species of *Saxicolella* occur mainly in the 700 – 1400 m altitudinal range, when other genera are predominantly restricted to lowland elevations. However, *Saxicolella nana, S. flabellata, S. deniseae* and *S*. sp. A all occur in the (100 –)400 – 700 m altitudinal band. The species of the genus appear to avoid coastal areas. Although Cameroon has the highest species diversity of both *Saxicolella* and Podostemaceae in Africa, *Saxicolella* is absent from the most species-diverse Podostemaceae site (which has 10 species) the Lobé Falls near Kribi, at the coast in the lowland evergreen forest belt (Cheek *et al*. 2017b). However, all but one of the eight species of *Saxicolella* co-occur at least once with one or several other species of Podostemaceae (see individual species accounts). The exception is *Saxicolella ijim*, which is was not observed to occur with other Podostemaceae, possibly because of its unusual ecological niche. *Saxicolella ijim* is unique in the genus in that it flowers in the spray-zone of a waterfall, and it is not immersed in water immediately before flowering as is usual in other species. However, *Ledermanniella prasina* J.J.Schenck & D.W.Thomas of the Korup has the same ecology (Schenk & Thomas 2004) and *L*.*letouzeyi* of the Bakossi Mts can also occur in the spray-zone of waterfalls although not exclusively as does *Saxicolella ijim* (Cheek *et al*. 2004).

### Pollination & Hybridisation

Although we suspect that pollination is by flying insects such as bees, as reported in other African podostemoids (Cheek *et al*. 2017b), no floral visitors have been reported or observed this far for any *Saxicolella* species. Hybridisation, reported for the first time in African Podostemaceae in Cheek *et al*. (2017b) is not known in *Saxicolella*. Since none of the species is sympatric, this is not unexpected.

### Habitat partitioning

In those four species of *Saxicolella* that co-occur at sites with other Podostemaceae species, it has not been possible to study habitat partitioning except for *Saxicolella futa* at one site in Guinea:

#### Case study: Salaa Falls, Futa Djalon, Guinea-Conakry

At this site four species of Podostemaceae occur in close proximity to each other, some tens of metres downstream from the main tourist falls. At one point, all four can be found within a 1m by 1m square. When observed by MC and DM in Jan. 2018, midway through the dry season, *Stonesia heterospathella* G. Taylor was in full flower, having been exposed by the slowly falling water in recent weeks, while *Ledermanniella guineense* C. Cusset growing deeper in the water than any other species, was just beginning to flower. Highest up the gradually sloping flat rock surfaces were colonies of *Tristicha trifaria* Spreng., long dead. *Saxicolella futa*, also long dead, grew 30 – 100 cm vertically above the water surface, in a band below the *Tristicha* and above the *Stonesia*, plants of two species intermingling at the interfaces. This same zonation, with *Tristicha* (above) and *Stonesia* (below) the *Saxicolella futa* was also seen just above the main Kambadga Falls near Pita, where the *Saxicolella* was much rarer.

The *Saxicolella futa* appears to grow, or compete better in slightly deeper water, than the *Tristicha*, and needs a shorter growing season (a shorter period underwater) than both the *Stonesia* and *Ledermanniella*.

**CONSERVATION STATUS**. The principal threats to *Saxicolella* species apply to Podostemoideae species as a whole, especially in Africa. Because they are restricted to habitats with clean, non-turbid, aerated water, with a rock substrate, degradation of any these environmental factors pose threats. Given that so many Podostemaceae species, including *Saxicolella*, are restricted to only one or two locations, they are especially at risk. A threat at even one location is likely to pose a high extinction risk for any *Saxicolella* present. All of the species are provisionally assessed as either Endangered or Critically Endangered using the IUCN 2012 standard, making this genus among the most threatened of its size globally.

### Turbidity & eutrophication threats

Turbidity in the water, indicating that silt is present, can reduce establishment of Podostemaceae seedlings (Philbrick & Novelo 1995). It can also reduce photosynthesis during the main growth period, when plants are under water in the wet season. (Cheek *et al*. 2015).

Algal growth can blanket Podostemaceae plants at some sites and reduce their ability to photosynthesise. Such growth appears to be associated with nutrient addition to rivers from human populations that may use water courses for processing crops, cleaning, and removal of waste-products. *Saxicolella futa* (this paper) appears to be threatened in this way.

### Hydroelectric Power Project threats

The greatest threats of global extinction for species of Podostemaceae such as those of the genus *Saxicolella* are from hydroelectric projects which have been growing rapidly in number in recent years as a source of cheap, greener energy in Africa. They are attractive to investors and governments being seen as sustainable and a good alternative to hydrocarbon-sourced energy. While hydroelectric projects have many environmental benefits compared with alternative options, all too often they threaten Podostemaceae species with extinction, and there are now many documented cases of local and global extinctions of Podostemaceae species resulting from such projects (Cheek *et al*. 2015; Cheek *et al*. 2017b; Couch *et al*. 2019).

Dams for hydro-electricity generation are constructed just above rapids or falls so as to benefit from the vertical drop in water levels (the “head”) at these sites. The construction of the dams may directly impact upon the falls and the species that they contain. More usually dams negatively affect populations of Podostemaceae through changes in water flow by four different threats:

1. Reduction of water flowing over falls at the dam site;
2. Impounding of water by the dam creates a large reservoir of motionless, non-aerated water in which Podostemaceae cannot survive;
3. Disruption downstream of natural season fluctuations in flow, important for Podostemaceae reproduction;
4. Cascade hydro projects which destroy all Podostemaceae habitat along the length of a river. These four threats are expanded in detail in Cheek *et al*. (2017b).

Cascade systems are steadily being developed in the Cuanza of Angola, where two of the four global locations of *Saxicolella angola* are thought to have been lost even before the species is formally named (see that species), in the Ogooué of Gabon which will threaten *S*. sp. A (see that species), and also in the Konkouré of Guinea which may already have destroyed the only known global population of *Saxicolella deniseae*.

### Difficulties with EIAs for Podostemaceae

It is extremely rare that competent Environmental Impact Studies (EIAs) are requested and conducted in advance of planning for such hydro projects in our experience. If EIA studies are conducted, they usually do not take into account the possible presence of Podostemaceae at these sites: many botanists mistake these flowering plants for mosses or algae (groups of plants usually regarded as non-threatened) and do not collect samples for identification so that dam construction goes ahead in ignorance of the presence of these often highly threatened species. Even if such studies have been done in advance, and samples collected from which Podostemaceae can be identified, two further obstacles exist: 1) many Podostemaceae have out-of-date Red List assessments which often misrepresent the species as being of low or unthreatened status when they may be highly threatened, and 2) most Podostemaceae species remain without a Red List assessment. Unless species can be shown to have a published Red List assessment of EN or CR on iucnredlist.org, or an extent of occurrence of <50,000 km^2^, they are generally not considered to merit concerted conservation action by the International Finance Corporation of the World Bank Group that often supports finance of such projects (IFC 2019).

**PHENOLOGY**. Species of the genus generally flower as water levels drop after the rainy season, exposing the plants that have developed underwater in previous months, and triggering flowering, and seed set, and if the plants become dried out, death. *Saxicola futa* is thought to complete its life-cycle in 6 months or less (see that species), but other species, such as *S. ijim*, may prove to be perennial if they are kept moist by waterfall spray throughout the dry season.

**ETYMOLOGY**. The name *Saxicolella* is compounded of saxicole, meaning ‘dweller on rock’ and – ella a diminuitive. The whole signifies “little dweller on rock” in Greek. However, Podostemaceae almost always grow on a rock substrate, and many are diminuitive.

**VERNACULAR NAMES**. None have been recorded. Generally, e.g. in Guinea, local communities do not have terms for different species of the genus at a location, but one term, treating the family as one entity (Cheek pers. obs, Guinea 2018, 2019).

### Generic delimitation & placement

*Saxicolella* has previously been used in a wider sense, to include superficially similar species (Cook & Rutishauser 2001; Ameka *et al*. 2002). However Koi *et al*.(2012) showed that, as a result, *Saxicolella* in that former, wider sense was polyphyletic. Re-evaluation of the morphology of the species revealed that two clearly defined genera are present, the second having the name *Pohliella* is characterised by bilocular ovaries, thread-like roots with haptera, distichous phyllotaxy, and rootcaps (see introduction and Table 1). For a synoptic revision of *Pohiella* see (Cheek 2020).

It can be argued that the three groups of species within *Saxicolella sensu stricto* could each be recognised as separate genera since in Asia, the same grouping characters (shoot position) have been found useful for this purpose (Koi *et al*. 2012, see above under shoots). Moreover, this argument is strengthened by the correlation of root characteristics with these same groups (see above) but correlated floral or fruit characters are lacking. Moreover, we are reluctant to increase the number of genera if there is an alternative option. Therefore, we have opted to recognise these three groups at subgeneric rather than generic level. Should molecular phylogenetic work support generic recognition (e.g., by long branches with high support values), consideration might then be given to elevate these subgenera to generic level. This would necessitate resurrection of the generic name *Butumia* G.Taylor (here adopted as a subgeneric name) and elevating the subgeneric name *Kinkonia* (proposed below for the eccentric *Saxicolella futa*) to generic level. However, it is possible that these groupings are the result of convergence and have no phylogenetic value. *Saxicolella sensu stricto*, sampled from Cameroonian material, is embedded within and near the base of the major clade of African podostemoids and is sister to all other African genera apart from *Inversodicraea* R.E.Fr. and *Monandriella* Engl. (Koi *et al*. 2012). The sister relationship of *Saxicolella* and *Monandriella* shown by (Koi *et al*. 2012) was foreshadowed by Engler who in his global treatment of Podostemaceae placed these two genera consecutively (Engler 1930: 29).

### Identification Key to the species of *Saxicolella*

1. Flowering shoots (0.9 –)4 – 10(– 21) cm long..... 2
1. Flowering shoots sessile or <0.5 cm long....... 4
2. Flowering shoots to 21 cm long; leaves dorsiventrally flattened, flabellate, bifurcating three times. Nigeria. ...........**1. *S. flabellata***
2. Flowering shoots to 7 cm long; leaves if flabellate, bifurcating only once. Cameroon and Angola ..............3
3. Flowering shoots (1 –)4 – 7 cm long, each with 4 – 6 spur branches; flowers single, terminating the short shoots. Cameroon. ......**2. *S. ijim***
3. Flowering shoots 0.9 – 1.5 cm long, unbranched; flowers in terminal cluster. Angola.............**3. S. *angola***
4. Roots 0.3 – 0.5(– 0.8) mm wide, bifurcating at intervals of 1.5 – 2.2 mm; shoots with spathellae single at the sinuses of bifurcations. ...**8. *S. futa***
4. Roots 1.8 – 4 mm (or more) wide, not, or rarely, bifurcating; shoots with spathellae closely spaced in centre of crustose roots OR at edge of ribbon-like roots but never at the bifurcations. .......5
5. Shoots several, stems distinctly visible, clustered in centre of the disc-shaped crustose root; distal leaves 1.5 – 6 mm long, divided 1.5 mm from the base into (2 –)3(– 4) filiform segments; .........**4. *S. nana***
5. Shoots without stems distinctly visible, in rows along the margins of the ribbon-like roots; distal leaves entire, not filiform; .............6
6. Leaves linear, flattened, exceeding the flower in length; pedicel at anthesis fully exposed, as long as ovary; ovary sessile (gynophore absent) ....**5. *S*. sp. A**
6. Leaves triangular or scale-like, far-shorter than the flower; pedicel at anthesis concealed inside the spathellum, far shorter than ovary..........7
7. Shoots with 5 – 7 ± isomorphic subulate leaves, which lack a large concave orbicular or elliptic basal part; leaves exceeding the ovary in length; stigmas complanate. W. Cameroon and S.E. Nigeria .........**6. *S. marginalis***
7. Shoots with 3 heteromorphic leaves, composed mainly of a concave orbicular or elliptic basal part; leaves all far shorter than the ovary; stigmas botuliform. Guinea .........**7. *S. deniseae***

**Subgenus 1. *Saxicolella***

Type species: *Saxicolella nana* Engl.

Roots approximately disc-like, crustose, with short radiating marginal lobes (where known). Shoots arising from the centre of the crustose root, forming distinctly visible stems with visible internodes; leaves with well-developed leaf-blades, far longer than the bases, usually bifurcating or trifurcating (usually entire in *S. ijim*), stipules present.

**ETYMOLOGY**. Autonym, taking the name of the genus.

**DISTRIBUTION**. Nigeria, Cameroon and Angola.

Species 1 – 4: *S. flabellata* (G. Taylor), C.Cusset, *S*.*ijim* Cheek, *S. angola* Cheek, *S. nana* Engl.

1. **Saxicolella flabellata** (G.Taylor) C.Cusset Cusset (1987: 94); Onana & Cheek (2011: 252 – 253); Onana (2011: 116); Onana (2013: 147). Type: Nigeria, Ogoja, Aboabom-Boje path crossing the Afi River, fl. fr. 13 Dec. 1950, *Keay* FHI 28240 (holotype BM, isotype K000435201) (Fig. 1). *Pohliella flabellata* G.Taylor (Taylor, 1953: 53; Taylor, in Keay 1954:124). Homotypic synonym.

**Fig. 1.**
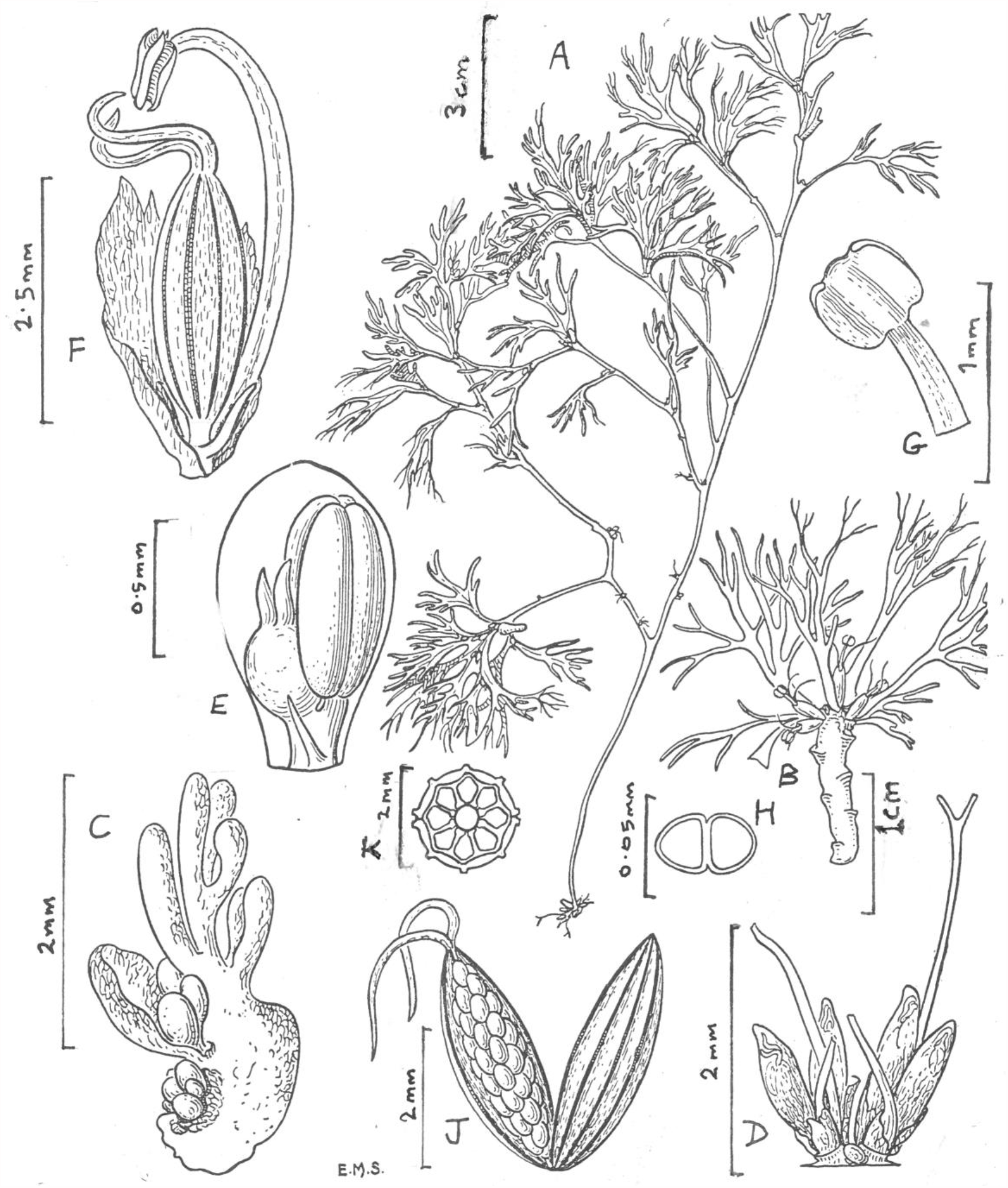
Saxicolella flabellata. **A** habit, flowering plant; **B** leafy, flowering shoot; **C** portion of root with young spathellae and shoots; **D** group of spathellae, each terminal from a root-borne leaf rosette; **E** developing flower inside spathellum; **F** mature flower (spathellum opened); **G** anther detail**; H** dyad pollen; **J** dehisced unilocular capsule showing seed; **K** transverse section of ovary (diagrammatic). From *Keay* in FHI 28240. ALL DRAWN BY E. MARGARET STONES (Originally published in Taylor (1953: 54) reproduced with permission of the estate of Margaret Stones and the Natural History Museum, London) © the estate of Margaret Stones.

*Perennial herb* (probably) with stems to c. 20 cm long, floating on surface of water when flowering. *Roots* incompletely known, recorded as green blotched red, ribbon-like, bearing both short shoots with sessile spathellae and long-stemmed shoots. *Stems* of long shoots terete, c. 2 mm diam., divided, internodes of principal axis 4.5 – 37 mm long. *Leaves* heteromorphic, leaves of long stems flabellate or dichotomously divided to 5 times, to 3 × 2 cm, ultimate segments capillary, base subpetiolate, sheathing, stipules absent. *Leaves* of short shoots root-borne, *s*ubtending spathellae 2 – 3, spirally inserted, outermost two leaves scale-like, sheathing, triangular-ovate or quadrate, slightly concave, 0.25 – 0.5 × 0.2 mm, second leaf longer than first; third leaf linear, 1 – 2.75 cm x 0.05 mm, apex obtuse, sometimes bifurcate. *Spathellae* in clusters of (1 –)2 – 4(– 5) either sessile on root or in leaf axils, cylindrical to narrowly ellipsoid (1.1 –)1.4 – 1.5 × 0.25 mm, apex obtuse, dehiscing at apex only. *Flower* at anthesis with ovary concealed within spathellum, only the styles and stamen exserted. *Pedicel* minute, 0 – 0.2 mm long. *Tepals* 2, subulate, 0.3 – 0.5 mm long. Staminal filament 4 – 4.5 mm long. Anther quadrate 0.5 – 0.75 × 0.5 mm, latrorse, pollen in dyads. Gynophore 0.2 – 0.25 mm long. Ovary ellipsoid to fusiform, 2.25 – 3.5 × 0.9 – 1 mm, with 8 longitudinal lines. Stigmas 2, filiform, 2 mm long. *Fruit* capsule ellipsoid, 3.5 × 0.75 mm, 8-ribbed, 2-valved. *Seeds* ellipsoid 0.25 × 0.15 mm (From Taylor, 1953 and *Keay* FHI 28240 (holotype K).

**RECOGNITION**. Distinct from all other known species of the genus in the very long stems, also in bearing spathellae from two different shoot type, those arising from the roots having different shaped leaves to those from the long stems. Similar to *Saxicolella angola* in the spathellae in clusters, and subtended by rosettes of more or less flabellate leaves (in other species the spathellae are single and leaves not remotely flabellate).

**DISTRIBUTION**. Nigeria, Cross River State, only known from the Afi River Forest Reserve near Ikom.

**SPECIMENS EXAMINED. NIGERIA**. Cross River State, Ogoja, Aboabom-Boje path crossing the Afi River, fl. fr. 13 Dec. 1950, *Keay* FHI 28240 (Holotype BM, isotype K000435201)(Only known from the type specimen.

**HABITAT**. River falls in evergreen forest, with *Ledermanniella tenuifolia* (G.Taylor)C.Cusset (G.Taylor 1954: 127 re *Keay* in FHI 28241). c. 113 m alt.

**CONSERVATION STATUS**. Ouedraogo (2010) assessed the conservation status of *Saxicolella flabellata* as Data Deficient in 2008, stating that there are records from Cameroon, Ghana, Niger and Nigeria. However, no records have been found from either Niger or Ghana and this seems most unlikely. Independently, on the basis of a location in Nigeria and one in Cameroon, Onana & Cheek (2011: 252 – 253) assessed the species as EN B2ab(iii). Kuetegue *et al*. (2019) also assess the species as EN B2ab(iii) citing no new data. Cusset (1987) had erroneously identified *Thomas* 2654 (K, MO, P, YA) collected 9 Dec. 1983 from Korup, Cameroon as this species, but this was corrected by Cheek (2020) to *Pohliella laciniata*. In fact, *Saxicolella flabellata* remains known only from the type collection made by Keay in Dec 1950 on a footpath across the Afi River in what is today the Afi River Forest Reserve of Nigeria. Reviewing Google Earth imagery for the site (placed at 6° 15’
s 28.6” N, 9° 00’ 42.71” E, elev. 113 m, viewed June 2021) shows that the footpath has been upgraded to a motor road, and that clearing of the forest canopy is steadily taking place confirming a report that dates from 6 Jan. 2016 and shows palm oil plantations, and open canopies indicative of logging, and these are confirmed by on-the-ground reports (Pandrillus). Surface run-off due to these activities may have contributed to extensive sediment deposits in the river viewed in the satellite imagery. Siltation of rivers is known to pose a threat to Podostemaceae (Cheek *et al*.2017b). Therefore, we assess *Saxicolella flabellata* here as Critically Endangered CR B1+B2ab(i-iv), estimating the AOO as 4 km^2^ as preferred by IUCN, and an EOO slightly larger.

**PHENOLOGY**. Flowering and fruiting in mid-December.

**ETYMOLOGY.** Referring to the shape of the leaves, flabellate meaning fan-shaped.

**VERNACULAR NAMES**. None are recorded.

**NOTES**. Originally described by Taylor as a *Pohliella*, he explained that he was in a quandary as to placement in this genus or in *Saxicolella* both of which were described by Engler (1926) which work he criticised (“I am not satisfied that the key characters used by Engler are sufficiently diagnostic”). In fact, Engler had separated these two genera in his key (Engler 1930: 29) based on locule number, and fruit rib number although they differ in other features. Taylor based his placement on features other than those in Engler’s key (“I have placed it in this genus due to the subulate stigmas and dichotomous leaves”), despite the first being discordant (“it deviates from the generic description in having a unilocular ovary”). It seems from the molecular phylogenetic evidence of Koi *et al*. (2012) that in this case locularity is indeed a better indication of relationships than leaf habit and stigma shape. *Saxicolella flabellata* is superficially very similar to *Pohliella laciniata* which grows in the same area: the Cameroon: Nigeria forest border. Both species flower from long stems when these are able to reach the water surface as the levels drop in the drier season. That both species have clusters of several flowers surrounded by rosettes of flabellate leaves that form a protective funnel, borne on long stems may be convergence to this scenario. These rosettes may function to float on the surface and protect the flowers they contain from water droplets (observed by the first author for *Pohliella laciniata* in Cameroon). The same trait (several flowers surrounded by a rosette of flabellate leaves) is seen otherwise seen only in *Saxicolella angola* where the ecology is unreported and the specimens fragmentary. Other species of *Saxicolella* have single flowers borne terminally in rosettes of leaves that are linear or highly reduced and can have no protective function during flowering (although they are likely to protect the developing flower buds).

Numerous other species are also both confined to the forest of the Afi River Forest Reserve and adjoining Cross River forests in S.E. Nigeria and known from only one or two collections, e.g., *Anchomanes nigritianus* Rendle (Moxon-Holt & Cheek 2021), *Talbotiella eketensis* Baker f. (Mackinder *et al*. 2010) and *Crossandra obanensis* Heine (Heine 1963).

2. **Saxicolella ijim** *Cheek* **sp. nov**. Type: Cameroon, NorthWest Region, Bamenda-Fundong, Anyajua, “Waterfall near Ijim Project HQ”, fl.fr., 12 Dec.1998, *Cheek et al*. 9920 (holotype K 000339632; isotypes SCA, YA). (Fig. 2) Syn. *Ledermanniella cf. musciformis sensu* Cheek (Cheek *et al*. 2000: 69,153)

**Fig. 2.**
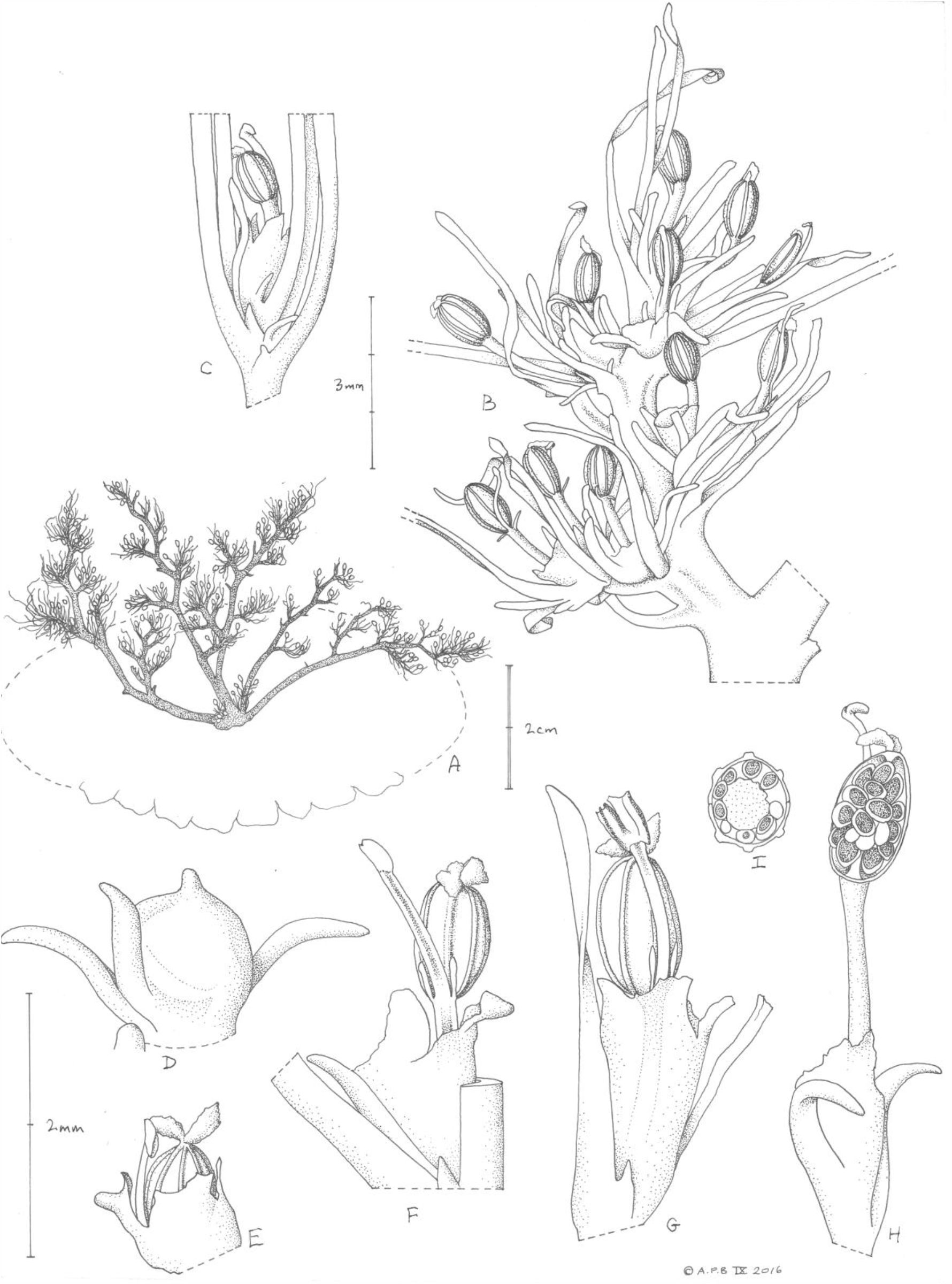
Saxicolella ijim. **A**. habit, showing crustose, disc-like root with radiating marginal lobes and centrally originating aerial stems; **B**. side-branch, fruiting; **C**. axillary, fruiting shoot; **D**. unopened spathellum; **E**. flower at anthesis, partly concealed in spathellum; **F**. & **G**. flowers at anthesis; **H**. fruiting shoot, one valve removed to show seeds on placenta; **I**. transverse section of fruit, showing absence of commissural ribs. From *Cheek et al*. 9920 (K). DRAWN BY ANDREW BROWN.

*Perennial or annual herb*, rosette-like, 7 – 8 cm diam. *Root* crustose in the central part of the plant, at the edge radiating and divided into separate, free, lobes, 0.5 – 0.8 cm wide (Fig. 2A). *Stems* (1 –)2 – 6, arising from the crustose centre of the root rosette (absent from the radiating lobes), erect, free-standing, branched from the base, (1 –)4 – 7 cm tall, spreading as wide as the root rosette, terete, each c. 2 mm diam. at base, with 4 – 8, ± evenly spaced, short leafy side-branches (Fig. 2B), proximal branches up to 1.5 cm long, phyllotaxy spiral, diam. slightly more slender than the principal axes, leaves with axillary rosette shoots. *Rosette (spur) shoots* axillary with stems inconspicuous, 1 – 3 per axil, each bearing 3 – 8 leaves and a single terminal spathellum. *Leaves of side branches* with spiral phyllotaxy, internodes c. 1mm long, laterally compressed, linear, (2 –)3.5 – 8 × 0.25 mm, entire, very rarely bifid, apex obtuse-rounded, basal 1 – 2 mm canaliculate, shortly sheathing the stem, astipulate (Fig. 2B), subtending axillary rosette shoots.

*Leaves of rosette shoots*, proximal leaves as those of the subtending side-branches, but usually with a pair of stipules arising from near base of the leaf sheath; distal 1 – 3 leaves immediately subtending the spathellum usually astipulate, shorter, 1.5 – 2 mm x 0.4 mm (Fig. 2D). Stipules symmetrical, equal, on each side of the leaf, narrowly triangular, 0.1 – 0.5(– 0.8) x 0.1mm, apex rounded, size of stipules increasing towards stem apex-spathellum, (Fig. 2C). *Spathellum* (undehisced) orbicular, 1 mm diam., mucro 0.2 mm long with apex rounded; dehiscing irregularly, post-dehiscence 1 – 2 × 0.7 – 1 mm (Fig. 2D). Flower ± erect in bud (in spathellum); at anthesis partly included in the ruptured spathellum (Fig. 2 E-G). *Pedicel* 0.5 – 1.5 mm long at anthesis. *Tepals* 2, slightly spatulate-oblanceolate to filiform 0.3 – 0.4 mm, distal portion 0.1 mm broad, flat, stipe 0.05 mm broad, erect (Fig. 2 F&G). *Stamen* as long as or exceeding gynoecium, filament 1 – 1.2 mm long, dorsiventrally flattened; anther oblong 0.5 × 0.25 mm. *Gynophore* 0(– 0.2) mm long. *Ovary* ellipsoid 0.75 – 1 × 0.6 – 0.65 × 0.7 – 0.75 mm, in transverse section slightly ellipsoid, slightly narrower along the sutured plane; unilocular, longitudinal ribs well-defined, 6 (three on each valve), commissural ribs absent (Fig. 2I). *Stigmas* 2, complanate, ovate, 0.25 – 0.4 × 0.18 – 0.2 mm, minutely verrucate (Fig. 2 E-H). *Fruit* about same size as ovary; pedicel accrescent (1.5 –)2.5 mm long, carrying fruit beyond the spathellum (Fig. 2H). *Seeds* ellipsoid 0.2 × 0.15 mm.

**RECOGNITION**. *Saxicolella ijim* differs from *Saxicolella flabellata* (G. Taylor) C. Cusset in the simple leaves, very rarely bifid (not flabellate, quadrifid); stigmas complanate, ovate (not filiform); fruit 6-ribbed, ellipsoid, length:breadth ratio c. 1: 0.65 (not 8-ribbed, fusiform, length: breadth c. 1: 0.33).

**DISTRIBUTION**. Cameroon, NorthWest Region, Bamenda-Fundong, Anyajua, known only from the type locality.

**SPECIMENS EXAMINED**. Cameroon, NorthWest Region, Bamenda-Fundong, Anyajua, “Waterfall near Ijim Project HQ”, fl. fr., 12 Dec.1998, *Cheek et al*. 9920 (holotype K 000339632; isotypes SCA, YA). Only the type specimen is known.

**HABITAT**. On boulders in spray zone below waterfall from basalt cliff, in former submontane forest belt. No other Podostemaceae present (Cheek pers. obs. Dec. 1998); 1300m alt.

**CONSERVATION STATUS**. *Saxicolella ijim* is known from a single waterfall, with only 30 – 40 plants scattered in an area of not more than 10 m x 10 m. Here the species is assessed as Critically Endangered, CR D. The waterfall is fed from a stream at the top of the Ijim Plateau where cattle have been introduced, posing a threat by their grazing and trampling increasing surface run-off and so silt levels in the stream feeding the falls. Targetted searches for Podostemaceae at numerous other waterfalls in the Fundong-Anyajua -Ijim area in 1998 did not uncover any additional sites for this species (Cheek *et al*. 1997; Cheek *et al*. 2000). Targetted searches by Ghogue to refind this taxon (then thought to be *Ledermanniella musciformis*) in the Bamenda area in 2006 with Ryoko Imachi and Yoko Kita failed to find it. Near-comprehensive botanical surveys in other locations S, W and E of Kilum-Ijim also failed to find additional locations although they brought to light several other species of Podostemaceae (e.g. Cable & Cheek 1998; Chapman & Chapman 2001; Harvey *et al*. 2004; Cheek *et al*. 2004; Cheek *et al*. 2010; Harvey *et al*. 2010; Cheek *et al*. 2011).

**PHENOLOGY**. Flowering and fruiting in Dec., 2 – 3 months after the end of the main wet season.

**ETYMOLOGY**. Named for Ijim, tribal lands of the Kom people, to which area this species is unique on current evidence.

**VERNACULAR NAMES**. None are known.

**NOTES**. When revisiting an incomplete and unsatisfactory identification the first author had made of a specimen from the Fondom of Kom in the Bamenda Highlands of Cameroon many years ago (*Cheek* 9920, Ijim, Anyajua, waterfall, 1300m, 12 Dec 1998, previously identified as *Ledermanniella* cf. *musciformis*: Cheek *et al*. 2000:153) it was realised that the fruiting ovary was erect emerging from the spathellum and, given the ribbon-like roots, longitudinally ribbed fruit and single stamen, that this could not possibly be a *Ledermanniella*, but a species of *Saxicolella*.

Prior to this paper, the only other published species of *Saxicolella sensu stricto* known which has long stems is *S. flabellata* but this new species differs in the shape of the leaves, ovary, and stigmas, and also in the number of fruit ribs and absence of a spathellum mucro. *Saxicolella ijim* occurs within a few kilometres of *S. marginalis*. These two species, together with *Saxicolella angola*, occur at the highest altitudes known for the genus (1300 – 1400 m alt.). The two Bamenda Highland species are easily separated since while the first has long stems, which are only produced from the centre of the radiating root rosette, the second lacks long stems completely, and instead bears numerous sessile, rosette-like stems along the margins of the radiating ribbon-like roots.

*Saxicolella ijim* is similar to *Saxicolella* sp. A in the unusual feature of the ovary being sessile (the gynophore being absent).

A summary of the rare, high altitude plant species of Kilum-Ijim (Mt Oku) is given by Maisels *et al*. (2000). Additional narrowly endemic species discovered from the Kilum-Ijim area are *Chassalia laikomensis* Cheek (Cheek & Csiba 2000), *Oxyanthus okuensis* Cheek & Sonké (Cheek & Sonké 2000), *Psychotria moseskemei* Cheek (Cheek & Csiba 2002), *Ternstroemia cameroonensis* Cheek (Cheek *et al*. 2017c), *Dovyalis cameroonensis* Cheek (Cheek & Ngolan 2007).

3. **Saxicolella angola** *Cheek* **sp. nov**. Type: Angola, Cuanza Sul, Gango-Cuanza, Mussende, 1000m alt., fl.fr. 31 June 1930 *Gossweiler* 9428 (holotype K000435202) (Fig. 3)

*Annual (probably) herb*, 1.2 – 1.9 cm tall. *Root* incompletely seen, probably crustose. Portion at base of stem (Fig. 3A) shield-like c. 1.1 mm diam., irregularly convex. *Stems* unbranched, erect, stout, self-supporting 9 – 15 mm long, terete, c. 0.5 mm wide at base, increasing to c. 1 mm diam. at apex. Proximal 1.8 – 5.5 mm lacking leaf-scars, distal portion with 5 – 7 leaf scars, internodes 0.5 mm long (proximal-most internodes) to 2 mm long (more distal internodes). Scars ± amplexicaul, distal nodes with leaf bases persistent, sheathing (Fig. 3B); stem apex with a head of flowers surrounded by clusters of heteromorphic leaves, phyllotaxy spiral. *Leaves* of outermost (proximal) part of apical cluster ligulate, 2.5 – 3 × 0.25 – 0.3 mm, or spatulate, that is with the distal end broader, elliptic, 0.8 × 0.4 mm, apex rounded or obtuse, base slightly sheathing, stipules absent (Fig. 3C); innermost, more distal leaves of apical cluster broadly ovate, or ovate in outline 0.6 – 1(– 1.5) x 1 – 1.5 mm, apex entire or slightly or deeply bifid, lobes equal or unequal, base broad, with or without marginal stipules. Stipules sometimes exceeding blades, subulate 0.5 × 0.2 mm. *Spathellae* in terminal cluster of (2 –)5 – 8, pre-dehiscence narrowly ellipsoid c. 2.5 × 0.8 mm, dehiscing into (2 –)3 – 5 subequal lobes, overall c. 2.5 × 1.5 mm. *Flower* partly concealed in spathellum at anthesis (Fig. 3D). Pedicel 0 – 0.5 mm long, concealed in spathellum. *Tepals* not seen. *Stamen* exceeding gynoecium, filament 2 – 2.05 mm long, anthers c. 0.4 × 0.3 mm. Pollen not seen. Gynophore 0.1 – 0.2 mm long. *Ovary* ellipsoid 1.5 – 1.8 × 0.6 mm, in transverse section suborbicular, unilocular; longitudinal ribs 6, ribs well-defined, 0.06 – 0.07 mm wide, commissural ribs absent. *Stigmas* 2, united at base, erect, cylindric, 0.4 – 0.45 × 0.1 mm, apex acute. Fruit about same size as ovary, mainly contained in spathellum (Fig. 3E) dehiscing by 2 equal valves, placenta spindle-shaped, 0.1 – 0.15 mm diam. Seeds ellipsoid c. 0.25 × 0.12 mm.

**Fig. 3.**
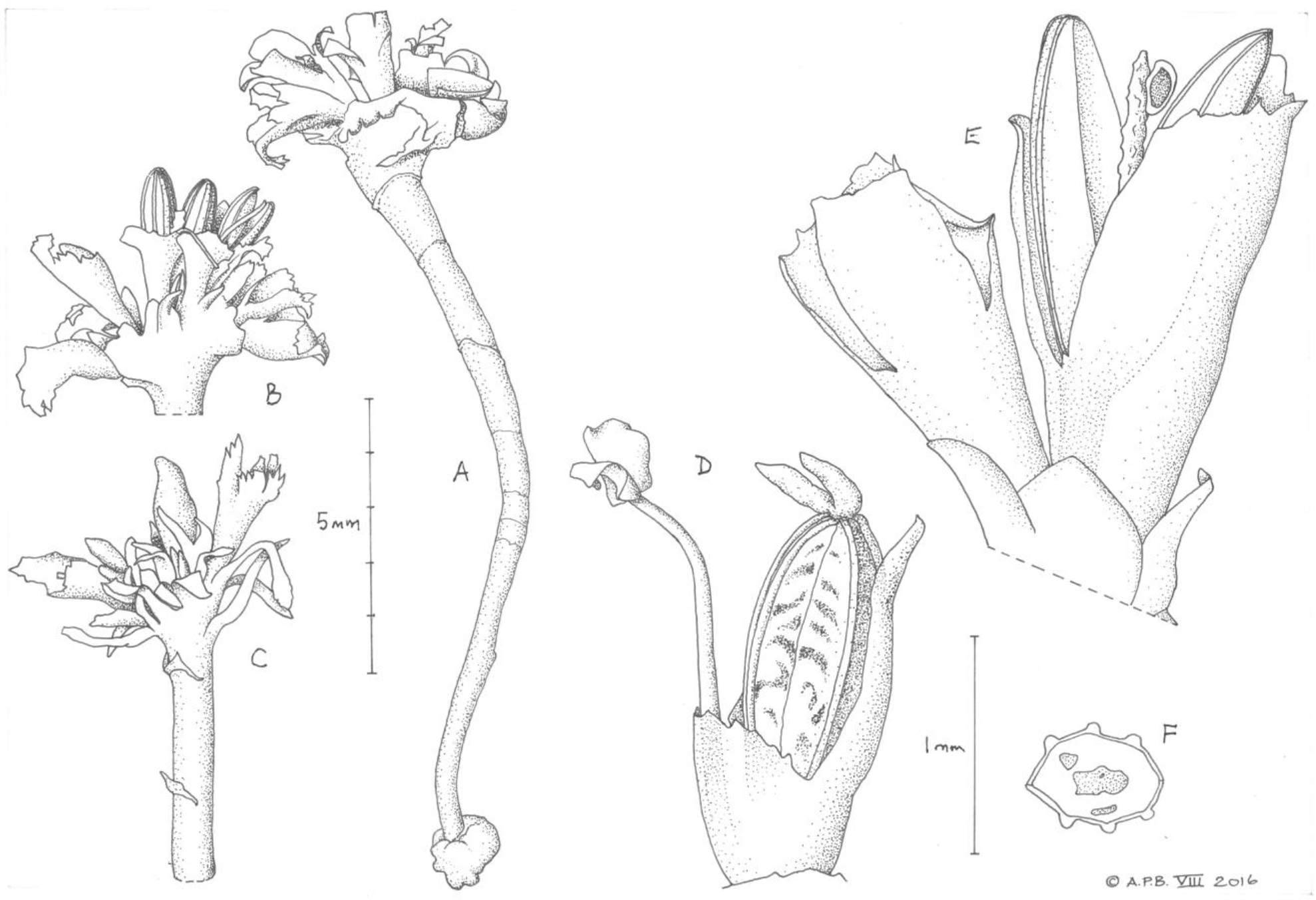
Saxicolella angola. **A** fruiting plant; **B, C** apices of two stems showing terminal clusters of leaves, spathellae and fruits; **D** flower, post-anthetic (anther empty); **E** two spathellae, one with dehisced fruit showing spindle-like placenta and a single seed attached; **F** transverse section of fruit (slightly distorted by compression). All from *Gossweiler* 9428 (Holotype K). DRAWN BY ANDREW BROWN.

**RECOGNITION**. Differs from other species of *Saxicolella* with elongated stems (*S*.*ijim* Cheek and *S. flabellata* (G. Taylor) C. Cusset in that the stems are unbranched (not highly branched), stigmas shortly cylindrical (not filiform, nor complanate); spathellae narrowly ellipsoid (not obovoid nor globose); leaves a mixture of simple-ligulate and ± isodiametric entire or bifid (not, only deeply quadrid, nor only simple-ligulate).

**DISTRIBUTION**. Angola, Cuanza Norte and Cuanza Sul Provinces (Cuanza River and its affluents).

**SPECIMENS EXAMINED**. ANGOLA, Cuanza Norte, Loanda Moaba, Duque da Bragança (now Kalandula Falls), Lucuala River, 1000m alt., “on rocks of river Loando near the waterfall”. fr. 29 August 1922, *Gossweiler* 8858 (K000593325); Cuanza Sul, Cuene or Cuno, bridge on river, “submersiherbosa, Limno Nereida” 650 m alt., 12 July 1937, *Gossweiler* 11353 (COI 00071845); Cuanza Norte, Cuanza River, Punta Filomeno da Camera, 100 m, submerged rocks, scarlet red plants, st. 7 March 1938, *Gossweiler* 12037 COI00071844

**HABITAT**. Waterfalls and rapids exposed to full sun, with gallery forest, 1000 – 1400 m alt. Occurring at the type locality near Mussende, with other species of Podostemaceae *Inversodicraea digitata* H.E. Hess and *Tristicha trifaria* (*Gossweiler* 9291, BM, n.v., COI00033957, ZT n.v.). At the Kalandula Falls, Cuanza Norte, occurring with *Ledermanniella aloides* (Engl.) C.Cusset (*Gossweiler* 8858A), *Tristicha trifaria* (*Gossweiler* 8858B) and *Inversodicraea fluitans* H.E. Hess (*Gossweiler* 8855, 8856, 8857) (Cheek *et al*. 2017b:139). At Punta Filomeno da Camera, occurring with an unidentified Podostemaceae collected later that year (25 June 1937) “in swift currents of water” (*Gossweiler* 10697 COI00033111).

**CONSERVATION STATUS**. Known from four collections, each at a different location of the Cuanza or an affluent. The collection site of “Punta Filomeno da Camera” has not been found but its altitude of 100 m on the Cuanza corresponds with the hydroelectric dam at Cambambe and so it is likely that the species has been lost at this location. However, the identification of the specimen is not completely certain since it was sterile and only viewed online. The site at “Cuene or Cuno, bridge on river”, since it is at 650 m alt. on the Cuanza, corresponds with the newly constructed Lauca Hydroelectric project, Angola’s largest. Here again the species is unlikely to survive due to the loss of its habitat and hydrological change. This leaves two locations upstream where the species is likely to survive. These two locations are c. 200 km apart from each other. *Gossweiler* 9428 (type specimen) near Mussende was recorded 31 June 1930. The exact site was not given, but is most likely to be the ford across the Gango River 16.5 km from Mussende on the road to Quibala (10°36’07.7” S, 15°52’49. 54” E observed on Google Earth), since this set of rapids is closest to Mussende (the town and river mentioned on the label) and is most readily accessible from that town. That vehicles are likely drive over the plants at the type locality is highly possible, and this would constitute a threat. The second locality, the Kalandula Falls (also known as the Calandula falls and formerly Duque de Bragança falls) on the Lucuala (or Licuala) River (*Gossweiler* 8858, 29 Aug. 1922), is now a major tourist attraction, probably because it is one of the major falls by volume in Africa and is only 360 km by road from the capital, Luanda. It is evident from the numerous posts of photographs by tourists on the internet (https://en.wikipedia.org/wiki/Kalandula_Falls accessed 30 May 2021) that trampling by visitors occurs, which can destroy plants of Podostemaceae as at the Lobe Falls in Cameroon (Cheek *et al*. 2017b). Both locations, are here ascribed an area of occupation of 4 km^2^ as preferred by IUCN (2012). Therefore, *Saxicolella angola* is here assessed as Endangered EN B2ab(iii) using the categories and criteria of IUCN (2012). Despite this assessment, and the fact that this species has not been recorded in the wild for 78 years, and despite the fact that Hess (1953) a Podostemaceae specialist who visited the Kalandula Falls in 1950 did not find the species there, we doubt that this species is extinct, although it is probably not common. This is because at both known locations, further apparently suitable habitat can be seen on Google Earth immediately upstream and/or downstream. However, there is no guarantee that the species occurs at these sites because it is rare and infrequent (only 2 of the 35 Angolan Podostemaceae held at COI are this taxon, and one of these is only doubtfully identified). Finally, there is no cause for complacency about the security of this species since the surviving two sites are both at risk of future new hydroelectric projects in which there is currently an upsurge in Africa and which are inimicable to the survival of Podostemaceae (Cheek *et al*. 2017b).

Angola is currently going through a surge of development posing risks to its species-diverse Flora. Additional range-restricted, newly described species endemic to Angola are *Barleria namba* I.Darbysh. (Darbyshire *et al*. 2019), *Barleria louiseana* I.Darbysh. (Darbyshire *et al*. 2021), *Justicia cubangensis* I.Darbysh. & Goyder (Darbyshire & Goyder 2019) and *Stomatanthes tundavalaensis* D.J.N. Hind (Hind & Goyder 2014).

**PHENOLOGY**. Only collected in flower (end June) and fruit (August) at the end of the dry season, the wet season being September to April

**ETYMOLOGY**. Named for the country of Angola as a noun in apposition. This species is both unique to Angola and the only species of the genus currently known to occur in the country.

**VERNACULAR NAMES**.

**NOTES**. *Saxicolella angola* was first recognised as a distinct species, but informally, and not published, by C. Cusset in 1975 and 1998. This is evident from her annotations of all four specimens of the species cited in this paper. She had annotated two specimens of what appears to be this taxon (*Gossweiler* 11353 and *Gossweiler* 12037 at COI, viewed online May 2021) as “*cf. Saxicolella angolensis*” dated 1975. Two other specimens, the basis of the description above, were sent on loan from K to P in 1982 (registered at K as H960/82) and were annotated as “*Saxicolella gossweileri* C. Cusset ined. 1998”, on one of which she had deleted an earlier annotation she had made of “*Pohliella angolensis* C.Cusset ined.”. Since the Code (Turland *et al*. 2018) advocates that such names should not be perpetuated without the permission of the author, and since that permission has not been obtained (attempts to contact her were unsuccessful), an alternative name has been selected.

*Saxicolella angola* is incompletely known. Both collections studied comprise of fruiting material. The roots, stem leaves, flowers at anthesis, pollen, are all either unknown, or only partially known. The ecology (microhabitat-ecological niche) is also unknown. Further field studies to fill these large gaps in our knowledge are advisable.

4. **Saxicolella nana** *Engl*. Engler (1926: 456 taf. XVII:1); Engler (1930: 48 fig 37); Onana & Cheek (2011: 254); Onana (2011: 116); Onana (2013: 147). Type: Cameroon, Centre Province, “In Nyong sudlich von Jaunde, Januar 1914” (in the Nyong S of Yaoundé, Jan. 1914) *Mildbraed* 7749a (holotype B10 0294988; isotype U1518023 n.v.)(Fig. 4)

*Annual herb* (probably), c. 5 mm high. *Root* crustose c. 5 mm diam., margin lobed. *Shoots* several from the centre of the crustose root, stems c. 1.5 mm long, bearing 3(– 4) spirally inserted leaves separated by short internodes, terminated by a single spathellum. *Leaves* 1.5 – 6 mm long, proximal part 1.5 – 1.6 mm terete c. 0.3 mm diam. distal (1 –)5 mm (2 –)3(– 4)-fid, the divisions equal, filiform. *Spathellum* pedunculate, peduncle to 2 mm long; spathellum obovoid-clavate, 2 – 3 × 0.9 mm, apex obtuse, opening by radial fissures, producing long triangular teeth. *Flower* carried c. 1 mm beyond the spathellum. Pedicel 2.5 – 3 mm long. Tepals filiform, 0.4 mm long. Stamen with filament 2 – 3 mm long, anther dimensions. Pollen in dyads. Gynophore 0.5 mm long. Ovary fusiform, 1.5 × 0.6 mm. Stigmas linear, 0.5 mm long. Fruit ellipsoid, dimensions as ovary, 6-ribbed, dehiscing by a single suture. (Description based on Engler 1926; Cusset 1987).

**Fig. 4.**
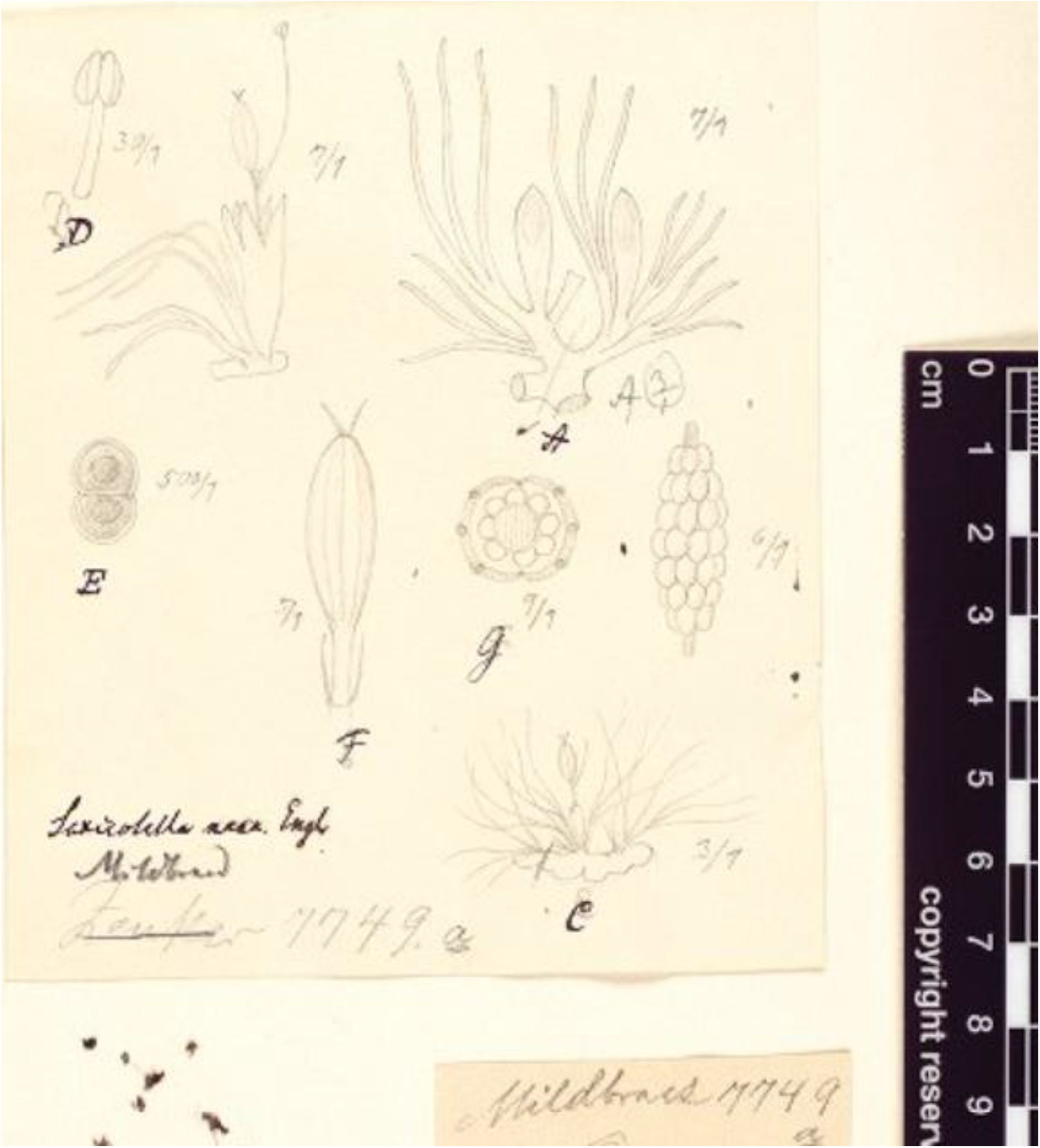
Saxicolella nana. **A** stem with two leafy shoots with spathellae; **B** placenta from fruit covered in seeds; **C** habit sketch showing disc-like root and centrally inserted leaf rosettes; **D** shoot with dehisced spathellum and flower at anthesis (right), detail of stamen (left); **E** dyad pollen grain; **F** flower showing pedicel, tepals and gynophore (stamen removed**); G** transverse section of ovary. All from *Mildbraed* 7749a (B). ALL DRAWN BY JOSEF POHL (original preparation drawing for illustration in the protologue, attached to the holotype B).

**RECOGNITION**. Similar to *Saxicolella* sp.A (see below) but differing in the disc-like root, the shoots produced at its centre (not ribbon like, the shoots at the margin), a distinct gynophore present, fruit 6-ribbed (versus ovary sessile, fruit 8-ribbed).

**DISTRIBUTION**. Cameroon, Centrale province. Only known from the type locality near Mbalmayo, Nyong River.

**SPECIMENS EXAMINED**. Cameroon, Centre Province, “In Nyong sudlich von Jaunde, Januar 1914” (in the Nyong S of Yaoundé, Jan. 1914) *Mildbraed* 7749a (holotype B10 0294988; isotype U1518023 n.v.); “11°27’N, 3°22’E” (likely 3°22’N, 11°27’E), 28 Feb. 2007, *M. Kato, R. Imaichi, S. Koi, & N. Katayama CMR-129* (Yan.v.) (cited in Kuetegue *et al*. (2019) see notes below).

**HABITAT**. Rapids in the river Nyong at Mbalmayo., in full light, in the semi-deciduous forest belt. Growing with *Macropodiella heteromorpha* (Baill.) C. Cusset (published as *Macropodiella mildbraedii* Engl. (Engler 1930: 48)). Alt. c. 330 m.

**CONSERVATION STATUS**. *Saxicolella nana* was assessed in 2007 as VU D2 (Ghogue 2010), citing a single location, the Nyong at Mbalmayo, with an AOO of <20 km^2^ and major threats of water pollution, temperature extremes and sudden drought. Independently, a provisional assessment of CR B2ab(iii) was made in Onana & Cheek (2011: 254), unaware that at this point, Ghogue had rediscovered the species likely at the type locality in 2004, the samples all being sent to Z. Cameroon has been relatively well-surveyed for Podostemaceae thanks to pioneering collectors in the German colonial period (1884 – 1916). More recently intensive surveys dedicated to finding sites for this family in Cameroon have been carried out above all by Ghogue, but also by dedicated Podostemaceae researchers from Switzerland, Ghana, USA, Japan and Britain but no further locations for *Saxicolella nana* have come to light. Kuetegue *et al*. (2019) studying the rheophytes of Cameroon concluded that it should be assessed as Endangered ENB2ab (ii, iii), with no additional data. Much of the length of the Nyong S of Yaoundé is not suitable for Podostemaceae due to the absence of rapids. However, some other rapids can be detected there on Google Earth (viewed June 2021) and would be worth visiting at the correct season to establish if the species has more than a single site. We know of no hydro-electric dams planned for the Nyong at present but this could well change given the number of dams planned elsewhere in Cameroon. We contend that the data presented merits reassessment as Critically Endangered CR B1+B2ab(i-iv).

**PHENOLOGY**. Flowering in January

**ETYMOLOGY**. The specific epithet refers to the small stature of the species.

**VERNACULAR NAMES**. None are known.

**NOTES**. The type at B was annotated by C. Cusset in 1974 as “*Pohliella nana* (Engl.) C. Cusset” suggesting that at that time she intended to make *Saxicolella* a synonym of *Pohliella*. She did not publish the combination and reversed the decision by the time she published her Flore du Cameroun account, when instead she sank *Pohliella* into *Saxicolella* (Cusset 1987).

The holotype at B bears three separate drawings. One of these appears to be made in the field by the collector, Mildbraed and shows the thalloid crustose root with a statement that the thallus firmly adheres for its entire surface. The second sketch (reproduced here as Fig 4) is the most comprehensive and shows numerous well-delineated thumbnail sketches of the dissected specimen some of which do not figure in the published illustrations such as Engler (1926), presumably due to space limitations. The third drawing is a more roughly drawn outline outline of parts of the second with an ink caption that may have been a guide to the journal designer in assembling the published image. The second drawing shows that both the stem and root thallus are better developed than that that is figured in the protologue, perhaps to save space. We credit the illustrations to Josef Pohl (for whom *Pohliella* was named) who provided scientific illustrations for Engler for 40 years (Anon. 2018)

Ghogue rediscovered *Saxicolella nana* in rapids in the Nyong near Mbalmayo, the presumed and likely type locality (details on the type label are indicative of this but not precise) which later in 2007 provided the source of material sequenced in phylogenies such as Koi *et al*. (2012). Such studies revealed that Engler (1926) was correct in erecting the two genera since *Saxicolella* in the broad sense as delimited by Cusset and followed by others, was shown to be polyphyletic (see introduction above). However, improved sampling of the species is desirable to confirm this conclusion.

When finalising this paper, recent records were found from Gabon on GBIF which are attributed to the species. However, from the images available, they seem to show several important differences from *Saxicollela nana* and are therefore provisionally identified in this paper as *Saxicolella* sp. A (see below under that species). If our hypothesis is incorrect and these specimens are shown to be *Saxicollela nana*, the extinction risk assessment for this species is likely to be reduced from the proposed CR to EN.

**Subgenus 2. *Butumia (****G. Taylor) Cheek stat. and comb. nov*.

Basionym: *Butumia* G. Taylor (1953:55)

Type species: *Saxicolella marginalis* (G. Taylor) Cheek.

Roots, crustose in centre, but mostly comprised of radiating, long, entire, rarely branched, broadly ribbon-like roots with numerous marginal shoots. Stems highly contracted, not visible. Leaves with sheathing base (where known), blade reduced, shorter than base (except *S*. sp. A), stipules present (were known).

**ETYMOLOGY**. Taking the name of the basionym *Butumia* (itself named for the Butum River, Nigeria).

**DISTRIBUTION**. Guinea-Conakry, Nigeria, Cameroon, Gabon. Species 5 – 7: *Saxicolella* sp. A., *S. marginalis* (G. Taylor) Cheek, *S. deniseae* Cheek,

5. **Saxicolella sp. A** Syn. *Saxicolella nana* auct. non Engl. (GBIF.org)

Prostrate herb with long, ribbon-like roots (black in life), infrequently bifurcating, Shoots sessile, forming 1-flowered leafy rosettes along margins of the root in rows, stem not visible. Leaves 3 – 6 per rosette, linear, flattened, as long or longer than the flower, entire, very rarely bifid at apex, apex acute to obtuse, stipules not seen. Spathellum caducous, sometimes visible as a broken ellipsoid remnant. Pedicel cylindrical, stout and white or pink in life, as long as ovary and nearly as broad. Stamen with filament 1.5 – 2 times as long as ovary, white to pink. Anthers with cells white, hemiglobose, isodiametric, semi-latrorse, each about as wide as filament. Gynophore absent. Tepals not seen. Ovary sessile narrowly ellipsoid, dull red, longitudinal lines not visible apart from commissure, orbicular in plan view (not laterally compressed). Styles 2, purple-red, erect stout cylindric, apices obtuse. Fruits 8-ribbed (*Texier et al*. 2321).

**RECOGNITION**. Differing from *Saxicolella nana* in that the root is long and ribbon-like with the shoots lacking visible stems, and arising along the margins of the root in rows (versus disc-like, crustose, the shoots with visible stems, arising from the centre of the root in a cluster); the leaves entire, linear, and not trifid from a point c. 1.5 mm from the base; the ovary sessile (the staminal filament inserted at its base), not with a distinct gynophore; fruit 8-ribbed (not 6-ribbed).

**DISTRIBUTION**. Gabon, Ogooué River and its affluent the Ivindo.

**SPECIMENS EXAMINED. GABON**. Ogooué-Ivindo Province, Ivindo National Park, Kongou waterfall area, islet in the middle of the Ivindou River, 00°17’32”N 012°35’23”E, 455 m alt. fl. 8 Feb. 2018, *Texier, Niangadouma & Akouangou* 2321 (BRLU, LBV, MO all n.v. photo); Haut-Ogooué Province, Boumango, Ogooué River, 02°16’27”S 013°38’20”E, 482 m alt., fl. 5 Aug. 2019, *Nguimbit* with *Boupoya, Ikabanga, Kaparidi* 26 (BRLU spirit only n.v.); Ogooué-Lolo Province, Chute Sessengué sur l’ Ogooué, 00°49’19”S 012°50’05”E, 323 m alt., fl. 31 July 2019, *Boupoya* with *Nguimbit, Kaparidi* 1952 (BRLU spirit only n.v., photo.).

**HABITAT**. Rocks in rivers sometimes (e.g. near Kongou Falls, *Boupoya* 1952) with other species of Podostemaceae, probably *Ledermanniella* (*Boupoya* 1953 – 1955, BRLU images); 323 – 482 m alt.

**CONSERVATION STATUS**. Once this species is formally published it will be possible for it to be Red Listed, most likely as Endangered EN B1ab(iii) since three locations are known with threats and extent of occurrence is calculated as 881 km^2^. Although the three points are widely separated they plot in a straight line forming a nearly linear polygon. *Texier* 2321 is stated to be “frequent on rocks on Ivindo River” but it is close to the Kongou falls. In 2007 the President of Gabon announced that a hydro-electric dam will be built at the Kongou falls inside the Ivindou National Park to support the Belinga Iron Ore project (Wikipedia, Kongou Falls, 2020). When implemented it can be expected to result in extinction of this species at this location as such projects have caused extinction of Podostemaceae elsewhere in Africa (Cheek *et al*. 2017b). While threats of this nature have not been found for the other two locations indicated, there is no scope for complacency. Six new large hydroelectric projects are planned in Gabon (Makoni 2020) and the total rises to 39 if potential and smaller projects are included (Cutler 2019). Apart from these, falls and rapids are the ideal sites for placing smaller hydro-power projects needed for future development projects. The location of *Nguimbit* 26 is close to an undeveloped iron ore project in Congo-Brazzaville, while *Boupoya* 1952 is about 10 km E of Lastoursville, one of Gabon’s major towns.

**PHENOLOGY**. Collected in flower in Feb. and July and August (dry seasons).

**VERNACULAR NAMES**. None are recorded.

**NOTES**. The first author came across this taxon when doing final checks for additional records of the genus on GBIF.org before finishing the manuscript for this paper. The three specimens were identified as *Saxicolella nana* respectively by Rutishauser, Bidault and Rutishauser, Bidault & Mesterhazy. An excellent image of *Texier et al*. 2321 and ten for *Boupoya* 1952 are available online through GBIF and Tropicos, (http://www.tropic.imageid=100597656 and http://www.tropic.imageid=100891363 respectively) but none for *Nguimbit* 26 which is included on the assumption that it matches the two specimens which had been identically named. Inspection of these images shows several characters that differ from those of *Saxicolella nana* which are given under “Recognition” above. No scale was available, so the description is made without giving dimensions, entirely on the basis of the online photos. Therefore, a full formal description of this taxon is still advisable, including observation to confirm the expected unilocularity of the ovary. It is possible that when the specimens themselves are studied the dissimilarities with *Saxicolella nana* will be reduced and that *Saxicolella* sp. A will be reduced to synonymy. Yet this seems highly unlikely given the number of qualitative morphological differences enumerated above. The specimens themselves could not be reviewed on loan because of technical difficulties due to their transfer from BRLU to ZT (Bidault pers. comm. to Cheek June 2021).

*Nguimbit* 26 was collected at a point so close (< 4 km) to the border with Congo-Brazzaville that it maps just inside that country on Tropicos. Since the river concerned extends far into that country it is very likely that the species occurs there and that it is not endemic to Gabon.

Gabon is seeing an upsurge in description of new species with 37 published in 2019, second only after Cameroon in tropical Africa, many of these being rare, range-restricted and endemic (Cheek *et al*. 2020b), joining those already documented narrow and threatened endemics as such *Diospyros rabiensis* Breteler (Breteler 1994), *Whitfieldia purpurata* Heine and *Whitfieldia rutilans* Heine (Grall & Darbyshire 2021), some of which are already extinct such as *Pseudohydrosme buettneri* Engler (Cheek *et al*. 2021a).

6. **Saxicolella marginalis** (G.Taylor) Cheek (Cheek *et al*. 2000:153; Onana & Cheek 2011: 253; Onana 2011:116; Onana 2013: 147) **Type:** Nigeria, “Ogoja Province, River Butum, Utanga, about 3 km north of Bagga, on smooth granite rocks, just below, at, and just above water-level in fast-flowing stream.”, fl. 25 Dec.1948, *Keay, Savory & Russell in FHI 25152* (Holotype BM000910416!)(Fig. 5).

**Fig. 5.**
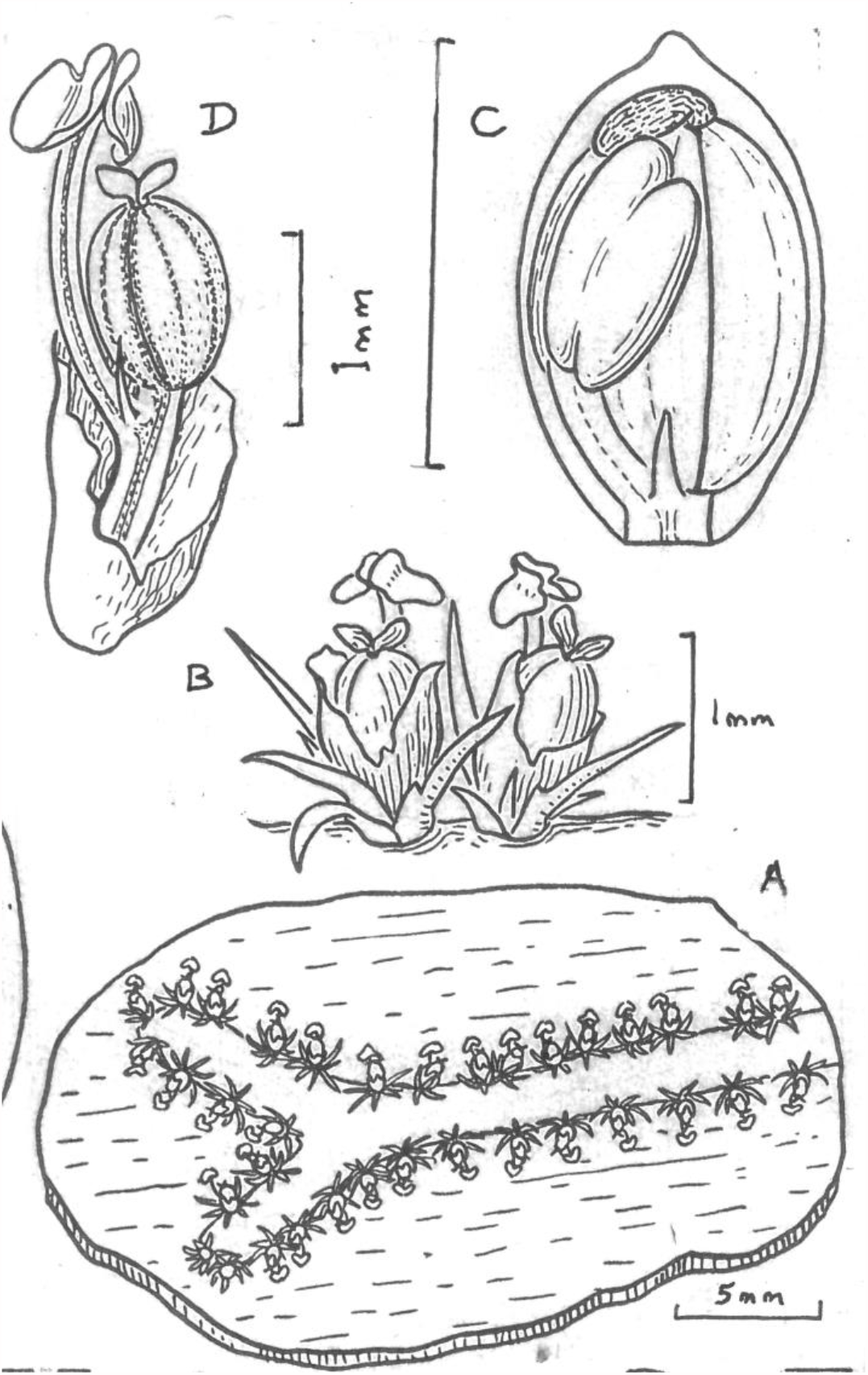
Saxicolella marginalis. **A** habit, flowering plant; **B** detail of flowering rosette shoots; **C** flower inside spathellum before anthesis; **D** flower at anthesis, spathellum opened. From *Keay* in FHI 25152. DRAWN BY MARGARET STONES. Originally published in Taylor (1954) as *Butumia marginalis* G.Taylor © the estate of Margaret Stones

*Butumia marginalis* G.Taylor (Taylor 1953: 55; Taylor 1954: 124 – 125). Homotypic synonym

*Annual herb*. Root rosette-like, 5 – 12 cm diam., radiating from a central point, the distal ends ribbon-like, 1.5 – 5(– 7.5) mm wide, rarely bifurcating. Stems of sessile leaf rosettes, numerous, spaced along the margins of the roots, (0.9 –)1.3 – 1.5(– 1.7) mm apart, 1.2 – 23 mm long (including the single terminal flowers), phyllotaxy spiral. Leaves (4 –)7 per stem, dimorphic, outermost (proximal), subulate, 0.25 mm long, lacking stipules; innermost (distal) 0.6 – 1(– 3) mm long, stipulate, stipules paired, marginal, triangular, 0.2 mm from base. Spathellum bud (pre-dehiscence) ellipsoid, 1.1 – 1.25 mm long, 0.6 mm wide, apex mucronate, dehiscing irregularly into two halves. Flower erect, included within spathellum at anthesis. Pedicel 0.9 – 1 mm long at anthesis. Tepals subulate, 0.4 – 0.5 mm long. Stamen exceeding gynoecium, 1.7 mm long, filament dorsiventrally flattened,1.2 mm long, anther oblong, 0.5 mm long, cells superposed. Gynophore 0.1 – 0.2 mm long. Ovary olive green, ellipsoid, 0.9 – 1 mm long, 0.6 – 0.75 mm wide, in transverse section orbicular, unilocular, longitudinal ribs 8, including two commissural ribs. Stigmas purple, 2, elliptic, complanate, 0.25 mm long, 0.15 mm wide. Fruit ellipsoid, 1 mm long, dehiscing into two persistent valves. Seeds ellipsoid, 0.2 mm long.

**DISTRIBUTION**. SE Nigeria and NW Region Cameroon, one site in each,

**SPECIMENS EXAMINED**. Cameroon, North West Region, Fundong, Touristic Hotel, 30 m fall on Chumni River, 1400 m, fl. fr. 22 Nov. 1996, *Cheek* 8740 (K!, SCA!, YA!)

**HABITAT**. waterfalls and rapids in forested or formerly forested mountainous areas, on rounded boulders of granite or basalt, in Nigeria with *Ledermanniella aloides* (Taylor 1954: 127 re *Keay* in FHI 25153); 400 – 1400 m alt.

**CONSERVATION STATUS**. Ouedraogo (2010) listed this species as Critically Endangered B1ab(iii)+2ab(iii), stating that it had only been recorded from Nigeria and Cameroon and that there is a continuing decline of its habitat quality due to water pollution and its populations are severely fragmented. He cited an earlier assessment of CR in 2000. This was probably that of Cheek in Cheek *et al*. (2000: 69) where the species was assessed as CR B1+2c due to the threat from laundry operations at the top of the fall housing this species in the town of Fundong, Cameroon.

Nevertheless, following the discovery of the species at the site in 1996, monitoring by the first author in December 1998 and November 1999 showed that approximately the same number of plants, several hundred, were present (Cheek *et al*. 2000: 69). This publication also formally transferred the species from *Butumia to Saxicolella*. Ouedrago (2010) further states that “the species may also be present in Ghana and Niger but this needs to be confirmed”. No evidence has been found to support this belief and it seems unlikely. Several other extremely rare, range-restricted and threatened species also occur at high altitude at Mt Oku (Maisels *at al*. 2000; Cheek *et al*. 1997; Cheek *et al*. 2000) e.g. *Kniphofia reflexa* Hutch. ex Codd (Hepper 1968), *Scleria cheekii* Bauters (Bauters *et al*. 2018), *Deinbollia onanae* Cheek (Cheek *et al*. 2021b)

*Saxicolella marginalis* is genuinely rare in Cameroon, since the first author has targeted searches for it at other falls and rapids in the Mt Oku area and not found it, although two other species of Podostemaceae have been found (Cheek *et al*. 2000). Near-comprehensive botanical surveys in other locations S, W and E of Kilum-Ijim have failed to find additional locations although they brought to light several other species of Podostemaceae (see references above under *Saxicolella ijim*). The status of the subpopulation at the type locality in Nigeria is unknown. The Red Data Book of Cameroon Plants (Onana & Cheek 2011: 253) reassess the species as EN B2ab(iii) since two locations (above) are recorded. Kuetegue *et al*. (2019) also assess the species as EN B2ab(iii) citing no new data.

**PHENOLOGY**. flowering and fruiting in late November to late December

**ETYMOLOGY**. the specific epithet derives from the rows of sessile shoots that line the margins of the roots.

**VERNACULAR NAMES**. None known.

**NOTES**. In describing this species as a new genus, *Butumia*, Taylor (1953: 57) stated: “Amongst African genera the plant is most closely related to *Saxicolella* and *Pohliella*, in each of which the flowers is unistaminate and erect within the spathella, but it differs from these genera in having entire rosulate leaves, much more shortly pedicellate flowers and complanate stigmas”. At that time both genera were only known from their type species and their circumscription was incompletely known.

It is remarkable that the only known Cameroonian site for this species is only a few kilometres distant and at similar altitude (c. 1400 m), to the only known site for *Saxicolella ijim* (see under that species). However, this submontane altitudinal band of the Cameroon Highlands is immensely rich rich in endemic range-restricted species. Other examples include *Coffea montekupensis* Stoffel. (Stoffelen *et al*. 1997) and *Impatiens etindensis* Cheek & Eb. Fischer (Cheek & Fischer 1999).

The Cameroonian location is about 100 km SSE of the type and only other known location in Nigeria. The species most closely similar in morphology to *Saxicolella marginalis* is *Saxicolella deniseae* of the Guinea Highlands in Guinea, far to the west. Both species share the unusual character of sessile, rosette-like shoots arranged along the margins of the radiating ribbon-like roots. Several species of the Cameroon Highlands do occur disjunctly in the highlands of Guinea e.g *Dorstenia astyanactis* Aké Assi (Couch *et al*. 2019), *Brachystephanus oreacanthus* Champl. (Champluvier & Darbyshire 2009) and *Isoglossa dispersa* Darbysh. & L. J. Pearce (Darbyshire & Pearce 2012), so long-distance dispersal, perhaps by birds, is credible as an explanation.

7. **Saxicolella densieae** *Cheek* **sp. nov**. Type: Guinea, Kindia-Télimelé Rd, 5 km S of crossing over the Konkouré River, on the Mayankouré near Lamba Sosso village, 10°24’15.4”N, 12°58’30.8”W, 180m alt. fl.fr. 27 Jan. 2018, *Molmou* 1683, with Gbamon Konomou (holotype HNG; isotype K000593321). (Fig. 6)

*Annual herb. Root* crustose, flat, covering the substrate in irregular oblong-shapes 5 – 18 × 7 – 24 cm, the margins with numerous radiating roots, each ribbon-like, 1.5 – 4 cm x 0.4 cm, tapering slightly to 0.25 cm wide towards the rounded apex (Fig. 6 A&B), not, or rarely, branching. *Shoots* sessile, short, monomorphic, at both margins of the root, rosettes (spiral phyllotaxy) of (2 –)3 leaves with a terminal spathellum, stems separated from each other by (0.15 –)0.18 – 0.25(– 0.4) cm. *Leaves* heteromorphic, proximal leaves suborbicular 0.3 – 0.5 cm diam., concave, sheathing, sometimes with a minute apiculus 0.05 – 0.1 cm long, stipules absent; distal leaves oblong-elliptic, 1.1 – 1.5 × 0.5(– 0.7) mm, including a short linear apical blade 0.3 – 0.6 × 0.1 – 0.15 mm, apex rounded; stipules equal, flanking the linear blade, triangular, c. 0.05 mm long, rarely absent. *Spathellum* pre-dehiscence ellipsoid c. 1.3 × 0.75 mm, mucro indistinct; at dehiscence c. 2.3 × 0.9 mm, with irregular dehiscence flaps. Flowers single per stem, terminal, erect in spathellum. Anthetic flowers with ovary half concealed inside spathellum (Fig. 6G). *Pedicel* erect 0.9 – 1 mm long at anthesis (0.15 mm long in undehisced spathellum) accrescent, increasing to 1.5 mm long in fruit. (Fig. 6J). *Tepals* 2, filiform, erect, 0.5 mm long, apex acute. *Stamen* 1, filament 1.6 – 3.6 mm long, slightly or far exceeding the ovary; anther cells oblong-ellipsoid, slightly diverging from each other in direction, (neither opposite, nor parallel) 0.25(– 0.4) mm long. *Gynoecium*, gynophore accrescent, 0.75 mm long in fruit. *Ovary* ellipsoid, at anthesis 1.2 × 0.4 mm, increasing in fruit to 2 × 0.65 mm, orbicular in transverse section, with eight more or less equal longitudinal ribs (commissural ribs well-developed). *Stigmas* 2 narrowly botuliform, 0.25 – 0.35 mm long, erect, minutely papillate, apices obtuse-acute. *Fruit* with placenta spindle-like. *Seeds* oblong-ellipsoid, 0.12 × 0.05 mm.

**Fig. 6.**
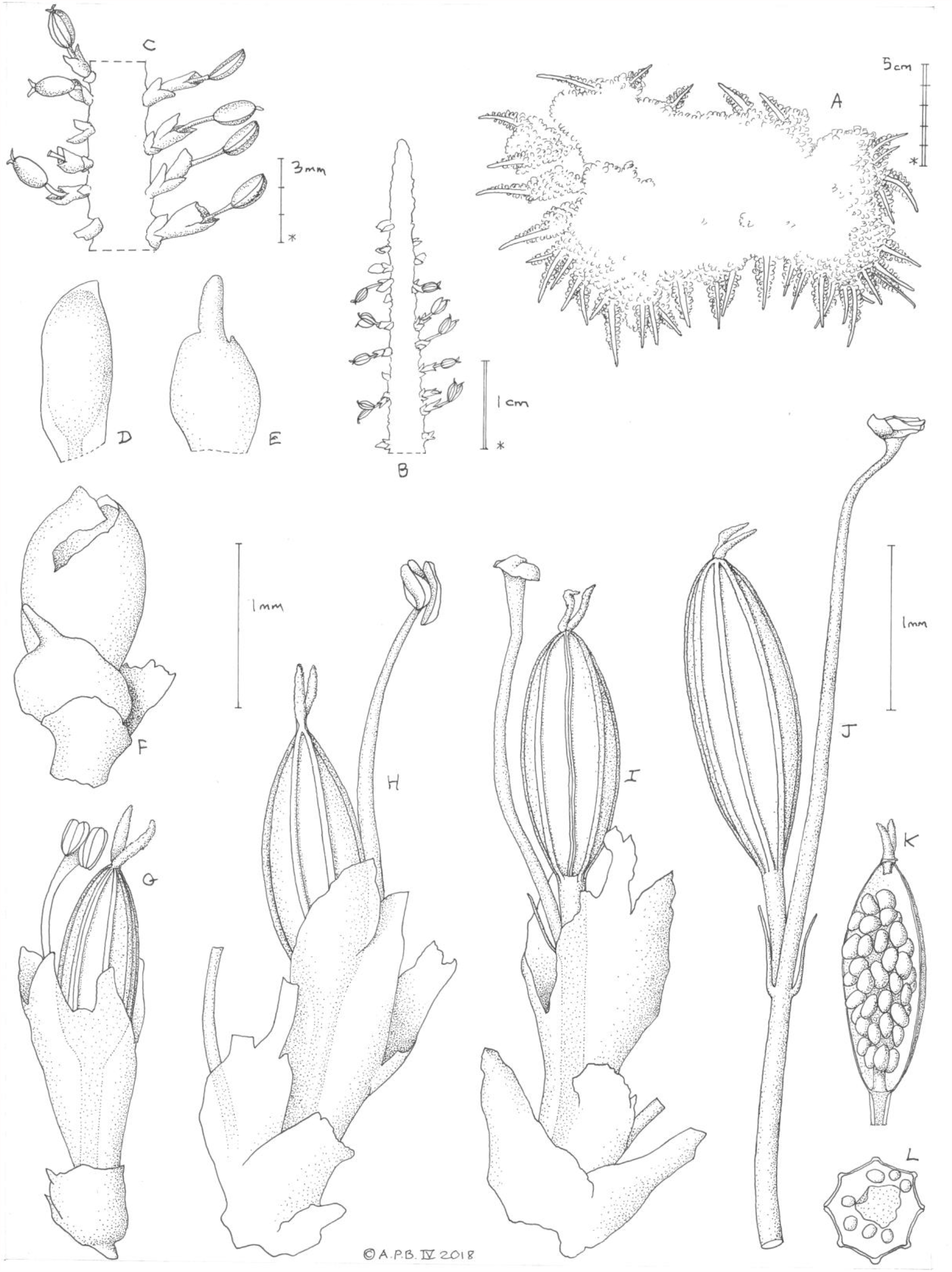
Saxicolella deniseae. **A** habit; **B** distal portion of one root, with marginal sessile shoots; **C** detail of B showing the fruiting shoots; **D** unopened spathellum; **E** distal leaf rib; **F** Shoot with opening spathellum; **G** & **H** shoots with flowers at anthesis; **I** & **J** fruit with persistent floral parts; **K** fruit with one valve removed showing seeds on the placenta; **L** fruit, transverse section. **A** from photo in habitat of and **B-L** from *Molmou* 1683. DRAWN BY ANDREW BROWN.

**RECOGNITION**. *Saxicolella deniseae* is most similar morphologically to *S. marginalis* (G. Taylor) Cheek because both species share numerous rosulate sessile stems inserted at the margin of the ribbon-like roots. *S. deniseae* differs in that the 3 rosette leaves are shorter than the ovary; and in that they are comprised of a concave orbicular sheathing base with only a rudimentary linear apical blade, the apex rounded (in *S. marginalis* there are 5 – 7 rosette leaves, each narrowly triangular-subulate, the sheathing base inconspicuous, the blade apex acute); in *S. deniseae* the stigmas are narrowly botuliform (in *S. marginalis* complanate).

**DISTRIBUTION**. Guinée, Kindia-Télimelé, Konkouré River.

**SPECIMENS EXAMINED**. Only known from the type specimen.

**HABITAT**. Rapids in river. Growing with several other Podostemaceae species, for which specimens for identification are not yet available. alt.174 m

**CONSERVATION STATUS**. Currently *Saxicolella deniseae* is known from a single location which will be impacted by a hydro-electric project in the near future (see notes below). The area of occupation is estimated as 4 km^2^ using the IUCN standard. A case of local extinction of Podostemaceae species due to hydroelectric projects in Africa is documented in Cheek *et al*. (2017b), and in Guinea in Cheek & Magassouba (2018) and in Couch *et al*. (2019). Therefore, we here assess *Saxicolella deniseae* as Critically Endangered (CR B2ab(i-iv)). It is to be hoped that further surveys will discover this species at other locations which are secure from threats such as hydroelectric projects otherwise this species is at high risk of extinction.

**PHENOLOGY**. Flowering and fruiting in January as the water level drops with the advance of the dry season. When collected in late January some plants were still emerging from the water, and live and flowering, while others had already been exposed and had concluded fruiting and died.

**ETYMOLOGY**. Named for Denise Molmou of Herbier National de Guinée, Université Gamal Abdel Nasser-Conakry, a leading botanist of Guinea and who led the botanical survey team in which this species was discovered and who also collected the type specimen.

**VERNACULAR NAMES**. None are known.

**NOTES**. Together with *Saxicolella futa*, this is the most westerly species of the genus. The two species are geographical outliers of the rest of the genus which is centred in the Cross-Sanaga interval of SE Nigeria and W Cameroon. *Saxicolella deniseae* cannot be confused with *S. futa* since in the first the ribbon-like roots are 0.4 cm wide, unbranched, and bear numerous sessile shoots along the root margin, while in *S. futa* the roots are 0.03 – 0.05(– 0.08) cm wide and bifurcate at regular intervals, bearing single sessile shoots only at the sinus of the bifurcations.

No associated endemic plant species are recorded near the location of *Saxicolella deniseae* because this part of Guinea is otherwise botanically unsurveyed.

### Hydroelectric dams on the Konkouré River of Guinea

*Saxicolella deniseae* is known from one site on the Mayan Kouré River, an affluent that, 5 km to the north joins the Konkouré, one of Guinea’s largest southward draining rivers. About 35 km upstream of that junction, at Donkheya, the Konkouré is dammed for a hydroelectric project. The reservoir extends upstream for 52 km (measured on Google Earth from imagery dated 31 Jan. 2015, downloaded 14 May 2018). This reservoir also extends up several of the affluents. About 30 km downstream of the same junction a new hydroelectric project was opened in 2017 (under construction in the Google Earth imagery of 31 Jan. 2015). The risk is that the reservoir of this project extends upstream a similar distance to that of Donkheya affecting the hydrology of its affluent, on the Mayan Kouré. Moreover, additional hydroelectric projects are planned on the Konkouré. The Konkouré river and its affluents remain almost completely unsurveyed for their plant species including Podostemaceae. Updating this before submission of this paper two years later, CNES Airbus imagery dated 11 Nov. 2018 viewed on Google Earth shows that as feared, the new reservoir on the Konkouré had already filled nearly up to the junction with its affluent, the Mayan Kouré. It is possible that by now, June 2021 the type location of *Saxicolella deniseae* has flooded which will have made the only known global population extinct.

**Subgenus 3. *Kinkonia*** *Cheek subgen. nov*.

Type species: *Saxicolella futa* Cheek sp. nov.

Roots radiating from a central point, lacking a central crustose part (or crustose portion minute), narrowly ribbon-like, bifurcating regularly and frequently, shoots only at the sinuses of the bifurcations. Stems highly contracted, inconspicuous. Leaves comprised of concave elliptic base, blade absent or rudimentary, stipules absent.

**ETYMOLOGY**. Named for the Kinkon Falls at Pita, one of the locations for the only known species.

**DISTRIBUTION**. Guinea-Conakry, Futa Djalon

Species 8: *Saxicolella futa*.

8. **Saxicolella futa** *Cheek* **sp. nov**. Type: Guinée (Republic of Guinea), Guinée Moyenne, Futa Djalon, Labé, Chutes de Salaa, 877 m alt., fr. 18 Jan. 2018. *Cheek* 18974 (holotype K000593322; isotype HNG). (Figs.7&8).

*Saxicolella futa* ined. Couch *et al*. (2019: 171, 214)

*Annual herb*, (1 –)1.5 – 3 cm diam., stems including fruits 1.8 – 2.2 mm tall (Fig.7A&B). Roots radiating, adhering strongly to substrate, dorsiventrally strongly flattened (thallus or ribbon-like), – 0.5(– 0.8) mm wide, internodes 1.5 – 2.2 mm long, repeatedly bifurcating at angles of 80 – 100(– 120°), drying bright white, resembling *Riccia* (thalloid liverwort) (Fig. 8). *Stems* monomorphic, extremely short, in the fork of root bifurcations, bearing a sessile rosette, phyllotaxy spiral, leaves 3 – 4, spathellum single, terminal. *Leaves* concave, irregularly ovate or elliptic, 0.3 – x 0.2 – 0.3 mm, apex acuminate, basal attachment broad, stipules absent, distal leaves larger than proximal. *Spathellum* pre-dehiscence ellipsoid, 0.4 × 0.25 mm, apiculate, basal part sheathed in leaves; dehiscing irregularly, then broadly funnel-shaped c. 1.2 × 0.8 mm. *Pedicel* erect, c. 0.6 mm long. *Tepals* 2, filiform, suberect, 0.2 mm long. *Stamen* one, filament 1.25 – 1.6 mm, erect, exceeding ovary; anther oblong, 0.18 – 0.2 × 0.18 mm, cells opposite. *Gynoecium*, gynophore curved, 0.6 mm x 0.1 mm. *Ovary* unilocular, ellipsoid in fruit, (0.7 –)0.8 – 0.85 × 0.4 – 0.5 mm, elliptic in transverse section, c. 0.4 × 0.5 mm (Fig. 7H), with six shallow longitudinal ribs (commissural ribs not developed). Stigmas 2, complanate, oblong, 0.175 – 0.2 × 0.1 mm. *Fruit* placenta spindle-like (Fig. 7E). *Seeds* oblong-ellipsoid 0.25 × 0.18 – 0.19 mm (hydrated) (Fig. 7J).

**Fig. 7.**
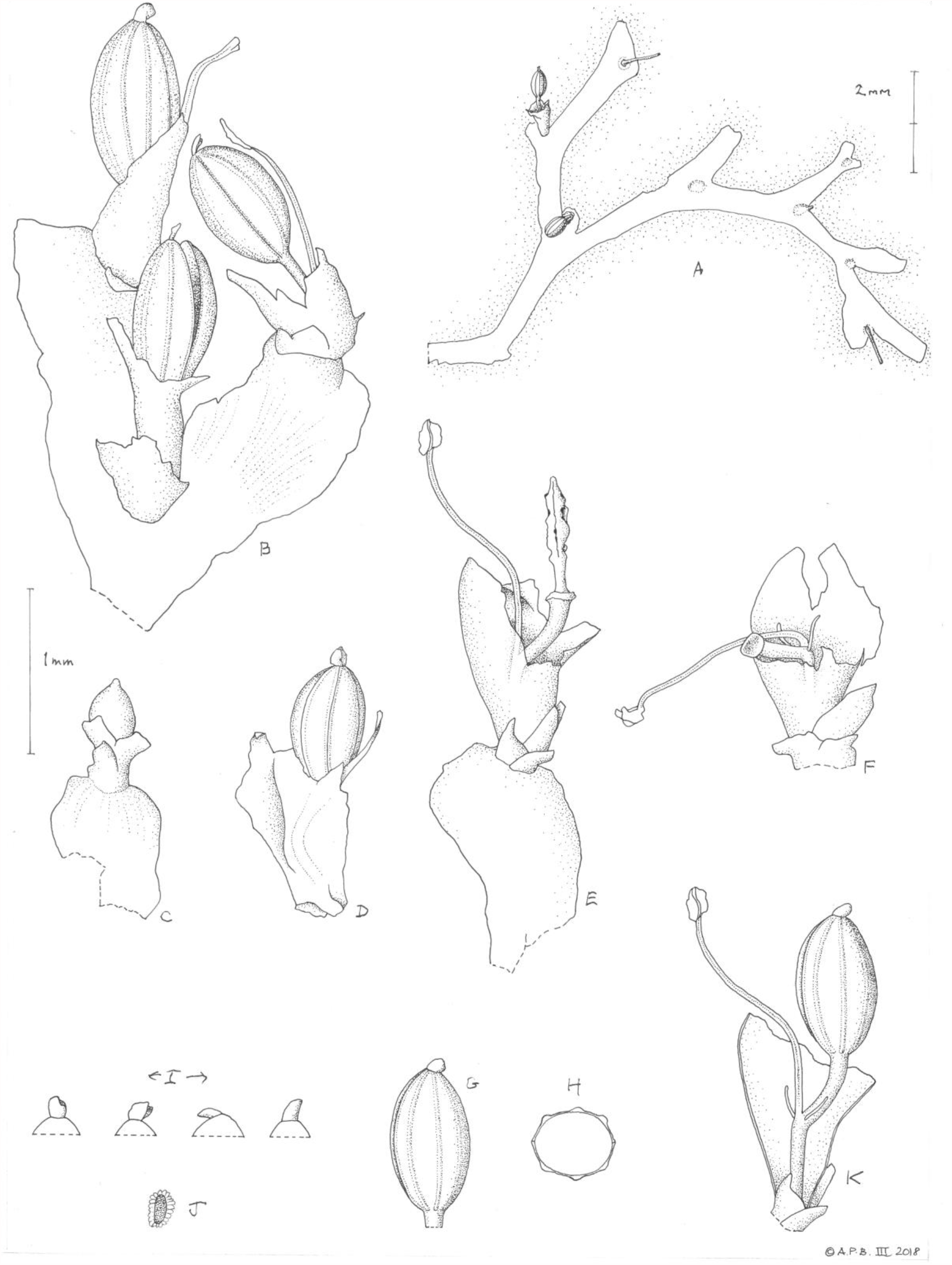
Saxicolella futa. **A** habit; **B** three flowering or fruiting shoots at root terminal-bifurcations; **C D** undehisced spathellum with shoot and root; **E** flower, part emerged from spathellum; **F** flower in spathellum, gynoeciou removed, showing tepals, gynophore and stamen; **G** gynoecium; **H** ovary wall in transverse section showing ribs; **I** variation in stigmata; **J** seed, hydrated; **K** reconstruction of complete flower (based on E-G). **A** from photo by Cheek at Chute de Sal’aa. **B-D, G-J** from *Cheek* 18979 (K); **E & F** from *Cheek* 18980 (K). DRAWN BY ANDREW BROWN.

**Fig. 8.**
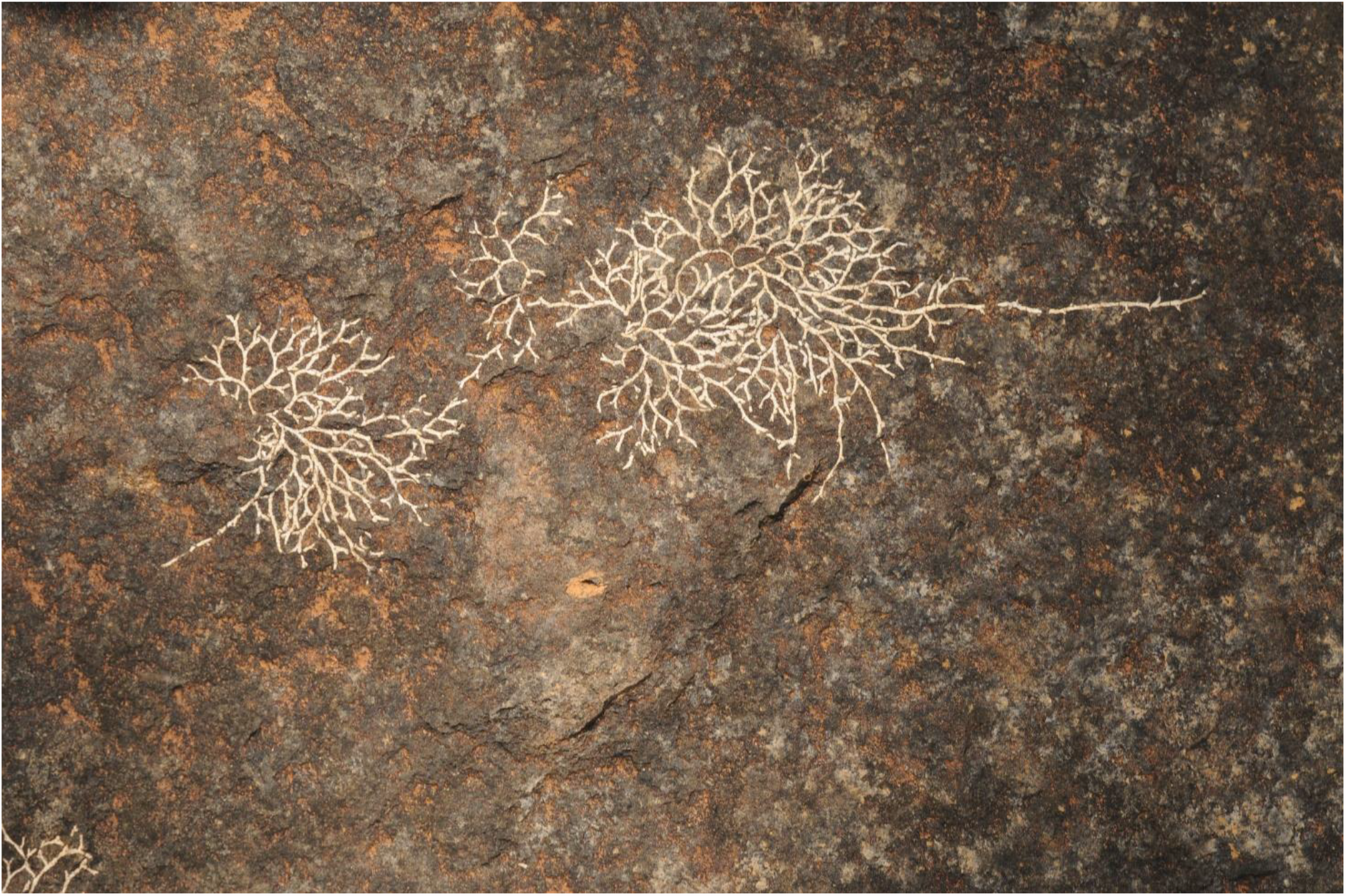
Saxicolella futa. Habit of the dead plant at the end of the dry season-start of the wet season. June 2016, Chute de Sal’aa. Photo. by M. Cheek.

**RECOGNITION**. Differing from other *Saxicolella* species with sessile spathellae (*S. marginalis, S. deniseae*) in the shoots not marginal, numerous, closedly spaced, but single, at root bifurcations; leaves lacking stipules (not stipulate); root slender (<1 mm wide, repeatedly and regularly bifurcating along its length (not >1 mm wide, not or rarely bifurcating along its length).

**DISTRIBUTION**. Guinée (Republic of Guinea), Guinée Moyenne, Futa Djalon.

**SPECIMENS EXAMINED**. GUINEA, Fouta Djalon, Pita, above Chutes de Kinkon, 857 m alt. fl. 18 Jan 2018, *Cheek* 18979 with Molmou (HNG, K000593323); ibid 18 Jan. 2018, *Cheek* 18980 with Molmou (HNG, K000593324); Pita, Chutes de Kambadga, 710 m alt. fr. 19 Jan. 2018, Cheek photo record; Labé, Chutes de Salaa, 877 m alt., fr. 18 Jan. 2018. *Cheek* 18974 (holotype K000593322; isotype HNG)

**HABITAT**. Waterfalls in the former cloud forest zone. The species grows at three sites each with several other Podostemaceae-Podostemoideae species (see case study above) but flowers, fruits and dies in advance of those, growing higher up in the riverbed than all other species of that group. 710 – 877 m alt.

**CONSERVATION STATUS**. Known so far from three locations: 1) Chutes de Salaa near Dalaba (type locality); 2) above the Chutes de Kinkon; 3) Chutes de Kambadga, downstream of Kinkon. The AOO is estimated as 12 km^2^ using the IUCN (2012)-preferred 2 km x 2 km gridcells and the extent of occurrence as 81.8 km^2^. Here *Saxicolella futa* is assessed as Endangered, EN B2ab(iii) since there are severe and imminent threats at all three locations. At locations 1) and 3) there are plans to build hydroelectric dams that are very likely to result in local extinction of the species (pers. comm. Cheek 2018). At site 2) the species is threatened by contamination of the water source by silt from run-off and by eutrophication due to contamination as the Kinkon river traverses the major town of Pita: less than 50 plants were seen at this location, and occupied a total area of < 3 m x 3 m. In the next 10 years this species is expected to be re-assessed as Critically Endangered (CR), or even “possibly extinct”. However, it is possible that further surveys may find additional sites for the species, which would be welcome. Seedbanking and public awareness actions will be put in place as soon as possible.

**PHENOLOGY**. Presumably germinating no earlier than May, with the beginning of the wet season, before which its habitat is dry. Flowering may begin as early as November after the conclusion of the wet season in October. By January fruit has formed and the plants are dead.

**ETYMOLOGY**. Named for the Futa Djalon highlands of the Republic of Guinea to which the species appears to be unique.

**VERNACULAR NAMES**. None are known.

**NOTES**. *Saxicolella futa* has a shorter growing season, (possibly completing its life-cycle in only six months or less) than all other Podostemaceae species present at each of the three locations at which it is known, apart from *Tristicha trifaria*. Both these two species were dead, dried and in fruit when encountered in Jan. 2018, while all other species of Podostemaceae present at these sites were still alive and only, for the most part, just becoming exposed by falling water and were in the process of beginning their flowering. The fact that, in these three locations, *Saxicolella futa* was found only on rock surfaces 30 – 100 cm above the water-level at which most other Podostemaceae (and all other Podostemoideae) occurred suggests that the species may have evolved into a niche to escape competition from those species. The extremely minute and inconspicuous stems, the diminutive ovaries, diminutive leaves, slender roots in comparison with other species of the genus, may well all be reductions that have enabled it to occupy a niche where the growing season is short, being the last to be submerged, and the first exposed, of all the podostemoideae niches present within its range.

*Saxicolella futa* is unique in the genus for its very slender (< 1 mm wide) ribbon-like roots, for the small size of its flowers, and for the position of the flowering shoots being only at the point of root. bifurcations, so that the roots appear to terminate in a shoot before bifurcating. It is also unusual in lacking evidence of stipules, but this is possibly concomitant with the reduction of leaf size maturity, more than 90 – 95% of the bulk of the plant consists of root, more so than any of the other species of the genus where the shoots are much more numerous and also larger and longer.

Recent surveys in Guinea connected with the Guinea Tropical Important Plant Area programme (TIPAs) have discovered several other new species to science, all of which are threatened. Several of these are like *Saxicolella futa*, also rheophytes, restricted to fast flowing water habitats, such as *Inversodicraea pepehabai* Cheek (Cheek & Haba 2016), *I. koukoutamba* and *I. tassing* (Cheek *et al*. 2019a), *Karima scarciesii* (Scott Elliot) Cheek (Cheek *et al*. 2016), *Lebbiea grandiflora* Cheek (Cheek & Lebbie 2018; Couch *et al*. 2019), while others are also found on the sandstone rock of the Futa such as *Keetia futa* (Cheek *et al*. 2018a), *Calophyllum africanum* (Cheek & Luke 2016), and *Kindia gangan* Cheek (Cheek *et al*. 2018b).

### Excluded species

*Saxicolella laciniata* (Engler) C.Cusset (Cusset 1987: 94) = *Pohliella laciniata* Engl.

*Saxicolella amicorum* J.B. Hall (Hall 1971: 133) = *Pohliella amicorum* (J.B. Hall) Cheek ined. (Cheek 2020)

*Saxicolella submersa* (J.B.Hall) C.D.K.Cook & Rutish. (Cook & Rutishauser 2001: 1165) = *Pohliella submersa* (J.B.Hall) Cheek

*Saxicolella macrothyrsa* A.Chev. (Chevalier 1938: 293) *nom illegit*.

## Discussion

The massive range extension of *Saxicolella* to the Guinea Highlands (due to recent collections representing new species to science) was unexpected, but recently has also been seen in other genera which were also considered previously to be confined to Lower Guinea (the region around the Bight of Biafra, far to the east). These genera are *Ternstroemia* Mutis ex L.f. and *Talbotiella* (Cheek *et al*. 2019b; van der Burgt *et al*. 2018, respectively). This is an indication of how incompletely surveyed the Guinea Highlands remain, despite already being known for endemic species and genera (Couch *et al*. 2019). New species are still being published regularly (Fischer *et al*. 2012; Phillipson *et al*. 2019; Cheek *et al*. 2020c).

In contrast, the range extension to Angola was not a surprise given that the first collection of the new taxon was made nearly a century ago, and its identification as a new species of *Saxicolella* more than 40 years ago. The delay in its formal publication speaks of both the shortage of taxonomists to do the work needed and the barriers to taking scientific discoveries through to formal publication. Despite the dramatic rise in numbers of the genus from three to eight, we still know very little of this genus in comparison with other African podostemoid genera. Additional species unknown to science of this and other Podostemaceae genera almost certainly remain to be discovered for science in Africa.

Until species are known to science, they cannot be assessed for their conservation status and the possibility of protecting them is minimal. About 2000 species of vascular plant have been described as new to science each year for the last decade or more (Cheek *et al*. 2020b). To maximise the survival prospects of range-restricted species such as Podostemaceae, there is an urgent need not only to document them formally but to assess them for their extinction risk, using the widely accepted IUCN Red List of Threatened Species (Bachman *et al*. 2019). Despite rapid increases over recent years in numbers of plant species represented by assessments on the Red List, the vast majority of plant species still lack such assessments (Nic Lughadha *et al*. 2020) and Podostemaceae are no exception.

As a global standard, the IUCN Red List supports the safeguarding and sustainability frameworks used by businesses and their major lenders (Bennun *et al*. 2018; Juffe-Bignoli *et al*. 2016). For example, clients of the International Finance Corporation (World Bank Group) are required to use the Red List to inform project risks and to refrain from activities leading to a net reduction in populations of species assessed on the Red List as Endangered (EN) or Critically Endangered (CR), over a reasonable timescale (IFC 2019). This is relevant because the International Finance Corporation have been linked with funding the hydro-electric projects that are major extinction risks for African Podostemaceae species.

Documented extinctions of plant species are increasing (Humphreys *et al*. 2019) and recent estimates suggest that as many as two fifths of the world’s plant species are now threatened with extinction (Nic Lughadha *et al*. 2020). That 100% of the species of a plant group with more than a few species should be highly threatened as documented in this paper for *Saxicolella* is unusual and may be unprecedented.

Extinctions of plant species are becoming documented throughout the range of *Saxicolella*. In Cameroon, centre of diversity of *Saxicolella*, the narrowly endemic species *Oxygyne triandra* Schltr. and *Afrothismia pachyantha* Schltr. are now known to be globally extinct (Cheek & Williams 1999, Cheek *et al*. 2018c, Cheek *et al*., 2019c) and *Vepris bali* Cheek was extinct even before it was known to science (Cheek *et al*. 2018d). Focussing on Podostemaceae, in Angola *Ledermanniella lunda* Cheek (Cheek *et al*. 2015) is thought to be globally extinct and in Guinea *Inversodicraea pygmaea* G.Taylor (Cheek & Magassouba 2018; Couch *et al*. 2019) both due to hydro-electric projects.

Efforts are now being made to delimit the highest priority areas in Cameroon for plant conservation as Tropical Important Plant Areas (TIPAs) using the revised IPA criteria set out in Darbyshire *et al*. (2017), as has already been completed for Guinea (Couch *et al*. 2019). These two countries have five of the eight species of *Saxicolella* between them and species such as *S. marginalis* and *S, ijim* (Cameroon) and *S. futa* (Guinea) are already included or set to be included in TIPAs. However, there is no such TIPA programme yet for Angola, Gabon and Nigeria, the other countries with species of *Saxicolella*. Even inclusion in an officially protected area is not a guarantee that a species will be safe from extinction. As in the case of *Saxicolella* sp. A of Gabon (see above), its most threatened location is inside a National Park, because a major Hydro-electric project is planned there. If new hydro-electric projects continue to be constructed there seems no doubt that global extinctions of Podostemaceae species will continue.

Alternatives to protecting threatened Podostemaceae species other than in their natural habitat are not yet available. While Podostemaceae have orthodox seed that can be banked, there are as yet, no documented cases of new populations being established artificially either experimentally or in the wild.

## Acknowledgements

The staff of the Université Gamal Abdel Nasser de Conakry-Herbier National de Guinée (UGAN-HNG), are thanked for arranging permits and facilitating the fieldwork that allowed the discovery of *Saxicolella futa* and *S. denisiae*. They are also thanked for long term support and collaboration under the Memorandum of Collaboration between UGAN-HNG and the Royal Botanic Gardens, Kew. Mr. Abdoulaye Yéro Baldé, Ministre, Guinean Ministère de l’Enseignement Supérieur et de la Recherche Scientifique, and Dr. Binko Mamady Touré, Secrétaire Général of the same Ministry, are thanked for co-operation. Colonel Namory Keita, Directeur, Direction National des Eaux et Forêts and Mr. Mamadou Bella Diallo, Point Focal CITES, Direction National des Eaux et Forêts authorised the export of the plant and seed specimens; permit numbers GN000053 and GN00018. This paper was initiated under the project “Important Plant Areas in the Republic of Guinea” supported by the Darwin Initiative of the Department of the Environment Food and Rural Affairs (DEFRA), UK government (project Ref. 23–002, 2016 –2019) by the Ellis Goodman Family Foundation (2018 – 2021), the Franklinia Foundation (2020 –) and the Critical Ecosystem Partnership Fund (2021 –). All supporting Kew’s Tropical Important Plant Areas (TIPAs) science strategic output partnership. Completion of work on the Cameroon species in this paper including *Saxicolella ijim* sp. nov. was completed through support for the Cameroon TIPAs programme from Players of the Peoples Lottery (PPL) and Red Listing work on the four Cameroon species is being supported by the John S Cohen Foundation.

John De Marco and Anne Gardner, then of BirdLife International’s Ijim Project and the Bamenda Highlands Forest Project based at Anajuya, North West Region, supported the discovery of *Saxicolella marginalis* in Cameroon, and (globally) of *S. ijim*. Sincere thanks to them both.

The then Heads of the National Herbarium Cameroon (YA), the late Dr Benoit Satabie and his successor Dr Gaston Achoundong, are thanked for expediting the partnership between Royal Botanic Gardens, Kew and the National Herbarium of Cameroon in 1996-1998 during which time the two collections above cited were made. Jean Michel Onana, Florence Ngo Ngwe, Eric Nana, and Jean Betti Lagarde, their successors, are thanked for support in maintaining this partnership.

Janis Shillito typed the manuscript. Eimear Nic Lughadha and two anonymous reviewers gave advice on earlier versions of the manuscript. Aurelie Grall kindly attempted to make contact with Colette Cusset concerning *Saxicolella angola*. Julia Buckley of the Department of Library, Art and Archives, Royal Botanic Gardens, Kew is thanked for arranging with the estate of Margaret Stones for permission to re-use Figures 1 and 5, which were originally labelled as *Pohliella flabellata Butumia marginalis* and first appeared in Taylor (1953 and 1954) respectively. Andrea Hart of the National History Museum also supported our re-use of Figure 1. Robert Vogt, Collections Manager B is thanked for expediting the loan to K of the type of *Saxicolella nana*.We thank Joaquim Santos, Collections Manager, Herbarium of the University of Coimbra, Portugal (COI) for help accessing the electronic collections databases of COI and LISC.

